# Cortex-wide laminar dynamics diverge during learning

**DOI:** 10.1101/2025.07.15.664840

**Authors:** Yael E. Pollak, Robert Sachdev, Matthew Larkum, Ariel Gilad

## Abstract

Learning to link sensory information to motor actions involves dynamic coordination across cortical layers and regions. However, the involvement of a particular layer in learning, especially from a cortex-wide perspective, is relatively unknown. Using wide-field calcium imaging in mice as they learn a whisker-based go/no-go task, we tracked activity in layers 2/3 or 5 (L2/3 or L5) across 25 cortical areas. A surprising initial effect of learning was that activity in L5 but not in L2/3 was globally suppressed at auditory cue onset. As the texture comes into touch, we found that L2/3 displayed learning-related enhancements in higher order association areas rostrolateral (RL) and secondary somatosensory (S2), whereas L5 in these areas oppositely decreased. During texture touch, the barrel cortex (BC) displayed similar learning-related enhancement in both layers. As sensory information is transformed into a motor action, there was a frontal/posterior divergence that emerges after learning, in which L5 was enhanced in the frontal cortex and L2/3 was suppressed in the posterior cortex. In general, learning related correlations were often stronger between distant cortical layers than within the same column, suggesting that learning drives laminar interactions that transcend traditional columnar organization. Together, these results reveal that learning orchestrates a dynamic interplay of activity across space, time and cortical layers. Our findings emphasize the critical role of laminar architecture in shaping cortical plasticity and support the view that layer-specific circuits are fundamental to sensorimotor learning.

## Introduction

Numerous forms of learning, memory and plasticity exist. A well-studied type of learning is associative learning, in which a stimulus is associated with a reward^1–6^. In associative learning paradigms, the subject is usually (or often) required to integrate an incoming sensory stimulus, associate it with a future outcome and act appropriately in order to receive the reward. Thus, learning a new task requires precise coordination and sensorimotor integration mediated by the neocortex^7–9^. Learning involves large-scale modulations across the cortex, including enhancement of relevant stimuli in primary sensory cortex^1,10–12^, dynamic reorganization and refinement of higher-order areas such as secondary somatosensory cortex (S2), association cortices^1,13^ and frontal motor cortices such as secondary motor cortex (M2)^14–17^. Learning-related modulations emerge during the association of sensory stimuli with actions and during the pre-stimulus period associated with cognitive functions such as attention and prior experience^18–20^. While much is known about how learning changes activity patterns in classes of neurons, very little is known about how the flow of activity changes across the brain for the two major classes of pyramidal neurons, the layer 2/3 (L2/3) and layer 5 (L5) neurons^1,11,21–23^.

Traditionally, L2/3 is considered the integrative layer both locally and between cortical areas, whereas L5 is considered the output layer, conveying information to subcortical areas along with motor information via the pyramidal tract^24^. But recent work shows that these distinctions are oversimplifications, L2/3 neurons across many cortical areas strongly encode movement^25–27^, L5 neurons encode stimulus properties and higher-order cognitive information^19,28,29^. In addition, the flow of activity can be local within a cortical column^30,31^ or global in which a specific layer in a specific cortical area project directly to another cortical layer in a different cortical area^32^. Locally, interactions between L2/3 and L5 can be excitatory or inhibitory, depend on the context, recording area and other parameters^33–36^. Globally, feedforward inputs, originating from lower-order sensory areas and the thalamus, ascend through the cortical hierarchy, targeting the middle layers, conveying external information relevant to the current task. In contrast, feedback signals, arising from higher-order or associative areas, descend to targeted superficial and deep layers, embedding prior knowledge, context, and predictions^30,31,37,38^. Two recent studies have addressed cortex-wide dynamics of different excitatory neuronal subtypes (primarily L2/3 vs. L5) and found diverse functionality across cortex especially in the frontal cortex^28,39^. These studies suggest that cortical decision-making emerges from the balance and integration of feedforward evidence, feedback modulation, and local recurrent computations within the laminar architecture.

Despite the extensive work on cortex-wide dynamics during learning, the specific contributions of neurons in individual layers are not known. Earlier work has shown that during motor learning, L5 neurons in primary motor cortex (M1) showed increased prediction accuracy, while L2/3 neurons maintained stable levels across learning^40^. In barrel cortex (BC), a subpopulation of L5 neurons became more responsive as mice learned to associate a certain stimulus with a reward^41^ and learning enhanced selectivity in L5 apical dendrites for both rewarded and non-rewarded whisker deflections^42,43^. Nevertheless, learning-related modulations in L5 neurons outside primary areas and at a more global scale are relatively unknown. In our previous work, we have imaged cortex wide L2/3 dynamics as mice use their whiskers to learn a go/no-go task^1^. We found a sequential increase in activity and correlation in BC and rostrolateral (RL) as a function of learning, whereas activity in other areas was suppressed. Here, we use a similar approach and compare cortex-wide L2/3 and L5 dynamics in 25 different cortical areas during learning. We find that some areas such as the BC display similar learning-related profiles in both layers, whereas other higher-order cortical areas including RL, S2 and posterior lateral cortex (PL) display opposing dynamics. In general, we find that as mice learn the task, L5 activity is globally suppressed at cue onset, followed by specific enhancements in L2/3 of task-related areas, which lead to widespread enhancements in L5 frontal cortex.

## Results

### Learning dynamics and behavioral parameters are similar in L2/3 and L5 transgenic mice

To compare learning-related dynamics at the cortex-wide level, we trained transgenic mice expressing GCaMP6f in L2/3 (Rasgrf-cre) or L5 (RBP4-cre) across the whole cortex on a whisker-based go/no-go texture discrimination task^1,18,26,44^. In short, two types of sandpaper were presented to the mice’s whiskers, and they were required to associate one of them (go texture) with a reward (Fig. 1a). In ‘Hit’ trials, the mice were rewarded for correctly licking in response to the go texture. They were punished with white noise for incorrectly licking in response to the no-go texture (‘false alarm’ trials, FA), and neither rewarded nor punished when they withheld licking for the go and no-go textures (‘Miss’ and ‘correct-rejection’, CR, trials, respectively). As mice learned to discriminate between the two textures, we used wide-field calcium imaging to measure cortex-wide neuronal population dynamics, in either L2/3 or L5^45^, along with concurrent video monitoring of whisking and body movements (Methods). In total, we imaged seven mice for each layer (L5 or L2/3) over 5–8 consecutive days (ranging from 1223 to 2494 trials per mouse).

**Fig. 1.**
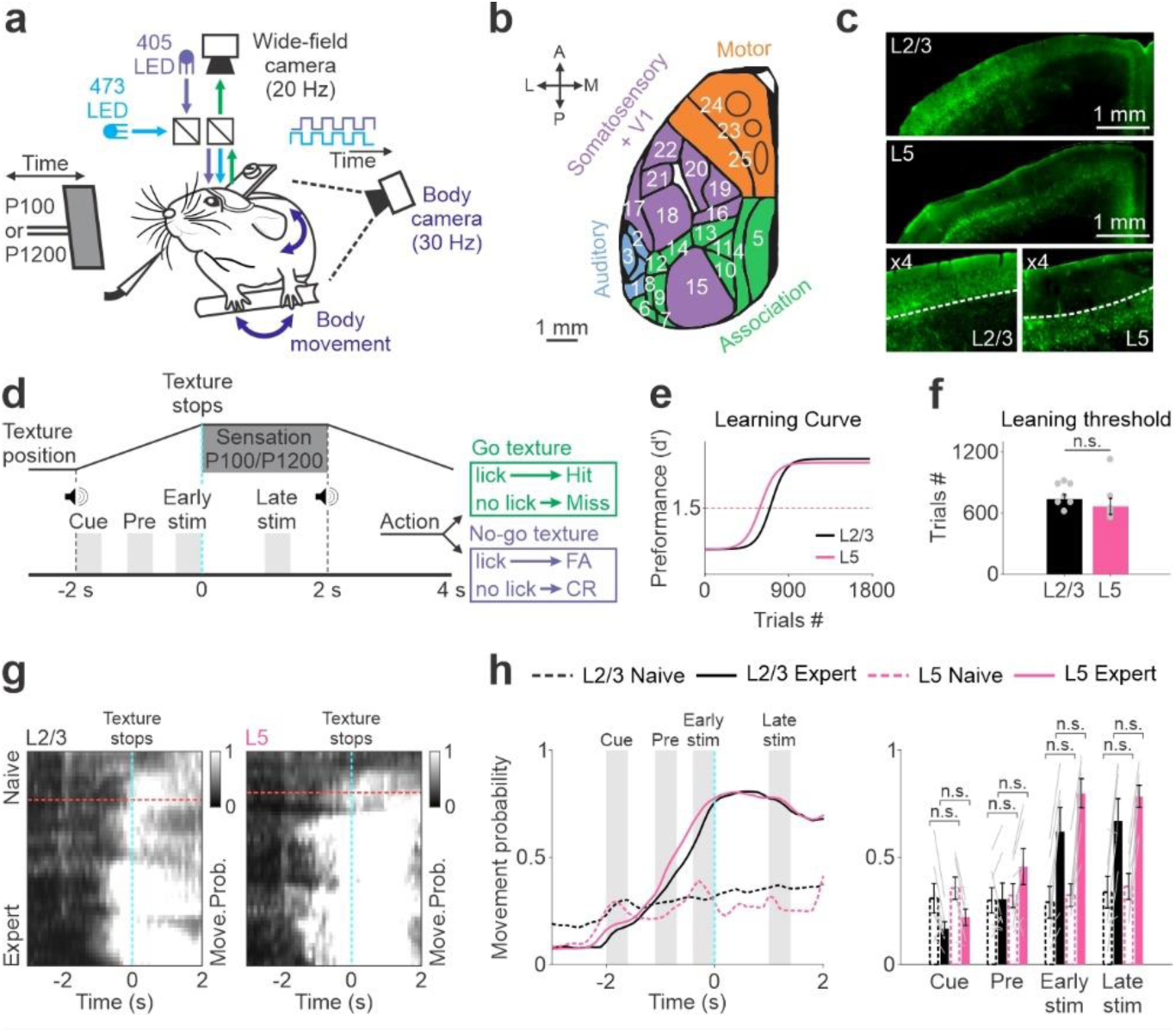
Mice groups of both L2/3 and L5 show similar learning profiles and movement strategies. **a** Schematic of Behavioral and imaging setup. **b** Top view of the dorsal cortex and 25 regions of interest 1. Temporal association (TEA) 2. Auditory dorsal (AD) 3. Auditory primary (A1) 4. Restrosplenial angular (RA) 5. Restrosplenial dorsal (RD) 6. Post rhinal (PR) 7. Posterior lateral (PL) 8. Lateral intermediate (LI) 9. Lateral medial (LM) 10. Posterior medial (PM) 11. Anterior medial (AM) 12. Anterior lateral (AL) 13. Anterior (A) 14. Rostro lateral (RL) 15. Primary visual (V1) 16. Primary trunk (Tr) 17. Secondary sensory (S2) 18. Barrel cortex (BC) 19. Primary hindlimb (HL) 20. Primary forelimb (FL) 21. Primary nose (No) 22. Primary mouth (Mo) 23. Whisker primary motor (M1) 24. Anterior lateral motor (ALM) 25. Secondary motor (M2) and four general divisions into auditory (blue), association (green), somatosensory + V1 (purple), and motor (orange). **c** Enlargement of a coronal slice displaying layer-specific (L2/3 or L5) green fluorescence. **d** Trial structure and possible trial outcomes. Distinct temporal periods are marked at the bottom (cue, pre, early-stim and late-stim). **e** Performance (d′) as a function of trial number averaged across mice of L2/3 (black; n = 7) and L5 (pink; n = 7; fitted with a sigmoid function). Red dashed horizontal line indicates threshold for learning (d′ = 1.5). **f** Learning threshold averaged across mice in L2/3 (black) and L5 (pink). Gray dots depict individual mice. **g** Movement probability for Hit trials of one example L2/3 (left) or L5 (right) plotted as heat maps along the two temporal dimensions (trial dimension on x-axis; learning dimension on y-axis; 30-trial bins along learning dimension). Horizontal red dashed line indicates learning threshold. Vertical cyan dashed lines depict the texture stop. **h** Left: Movement probability along the trial averaged across L2/3 (black) and L5 (pink) mice during in naïve (dashed lines) and expert (solid lines) cases. Gray bars indicate temporal periods. Right: Movement probability averaged across L2/3 or L5 mice within each temporal period. Error bars depict mean ± s.e.m across mice. Gray lines depict individual mice. n.s. not significant, Rank-sum test.

Cortex-wide dynamics were aligned together across days and registered onto the 2D top view Allen reference atlas, 25 cortical areas were defined and divided into four general groups^1,18,46,47^ (Motor, Somatosensory + V1, Association and Auditory, Fig. 1b; Methods^1^). The trial was initiated with an auditory cue, which signaled the approach of a texture within whisker touch for a duration of 2 seconds. Then, the texture moved out of position and a report window of 2 seconds started (Fig. 1d). We defined four discrete time periods in the trial: the ‘cue-period’, which began just after the auditory cue associated with the texture starting to move towards the whiskers (2 to 0.7 s before the texture stop); the ‘pre-period’ (1.1 to 0.7 s before the texture stopped) as the texture approached the whiskers; the ‘early-stim period’ (0.4 to 0 s before the texture stopped), which includes the epoch when the whiskers touched the texture, and the ‘late-stim period’ (1 to 1.4 s after the texture stopped) when the mouse transforms its sensation into an action plan (in Hit trials). The trial structure remained constant across learning.

All mice increased performance with training (5–8 days; ∼400 trials/day) and eventually reached expert performance (d′ within 30-trial bins; Fig. 1e and Supplementary Fig. 1a for individual mice; see Methods). Importantly, the learning profile of L2/3 and L5 mice are similar (Fig. 1e; Supplementary Fig. 1a for individual mice). For each mouse, we defined the ‘learning threshold’ by crossing a d′ of 1.5, and the ‘naïve’ and ‘expert’ phases as 300 trials, approximately 150 trials before and after the crossing point, respectively. The learning threshold is not significantly different between L2/3 and L5 mice (p > 0.05; Rank-sum test; Fig. 1f).

Next, we quantified the body movement of the mice as a function of learning, which was shown to have a profound effect on cortex-wide neuronal dynamics^1,25–27^. To do this, for each trial, we extracted forelimb and back movements from the body camera and calculated a binary movement vector by choosing a fixed threshold^1,26^ (30-trial bins; Methods). An example of a 2D movement probability (only Hit trials across learning) reveals that as mice gained expertise, they exert increased body movement, which initiated earlier, as texture approach the whiskers (Fig. 1g). Importantly, L2/3 and L5 mice exhibit insignificant differences in movement probability along the trial in both naïve and expert cases (Fig. 1h; p > 0.05; Rank-sum test; Supplementary Fig. 1b for individual). In summary, L5 and L2/3 mice display similar learning profiles along with similar movement strategies, enabling us to compare learning-related cortex-wide dynamics between layers.

### Learning-related laminar divergence during sensorimotor integration

We analyzed spatiotemporal dynamics of L2/3 and L5 cortical activity across learning, as revealed by wide-field calcium imaging. Here, we focused on Hit trials, in which probabilities were relatively high across learning and similar across layers. In general, both L2/3 and L5 display similar temporal dynamics during the trial (Fig. 2a), in which RL association cortex is activated as the texture approaches the whiskers and precede the BC, which is mainly activated during the texture touch (early-stim period^1^). When comparing activity maps between naïve and expert mice within a specific layer, we observe several distinguishing features within each time period across L2/3 and L5. For example, during the cue-period, L5 naïve activity map shows widespread responses that are less evident in the expert map. This change is not evident in the L2/3 maps.

**Fig. 2.**
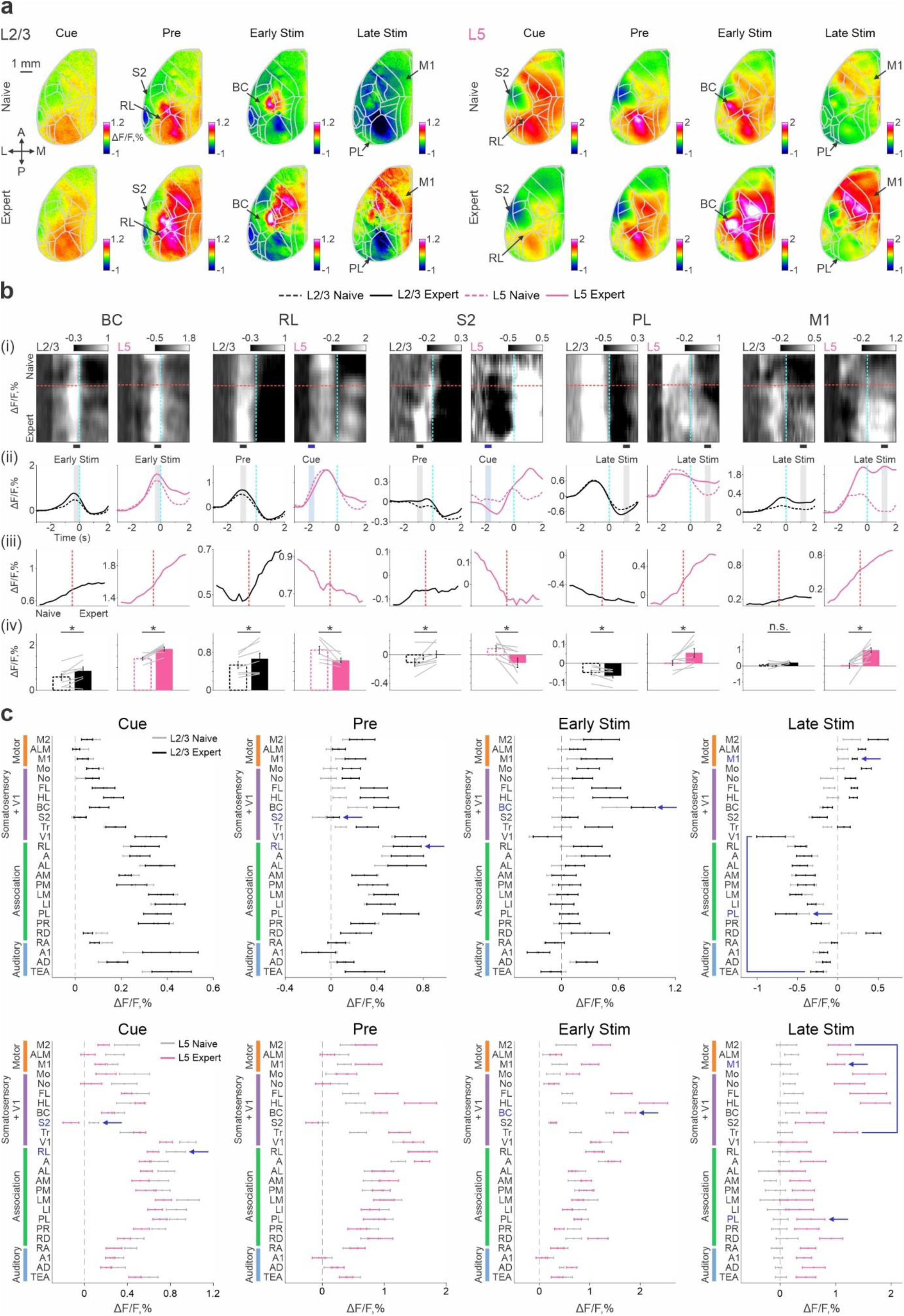
Cortex-wide laminar differences during learning. **a** Example activation maps from one mouse of L2/3 (left) and L5 (right) averaged during cue, pre, early-stim and late-stim periods in naïve (top) and expert (bottom) phase. 5 areas of interests are marked. Color scale bar indicates min/max of percent ΔF/F. Overlay of areas in gray for all maps. Scale bar is16.02 cm 1 mm. **b** (i) Example of a 2D heat map of response (ΔF/F) from one L2/3 (left) and L5 (right) mouse (trial structure along x-axis; learning scale along y-axis) for five areas of interest (from left to right): BC, RL, S2, PL, and M1. Data is binned every 30 trials. Dashed cyan line indicates texture stop. Dashed red line indicates learning threshold. (ii) Temporal response along the trial for each area averaged across L2/3 (black; n = 7) and L5 (pink; n = 7) mice for expert (solid line) and naïve (dashed line). Time period for further analysis window is marked by gray or blue colors. (iii) Mean responses across learning for each area averaged during the specific time period indicated in the plot above (n = 7 mice for each layer). (iv) Mean response for each area during the specific time period in expert (straight line) and naïve (dashed line) cases. Error bars are s.e.m. across mice (n = 7). Lines depict individual mice. *p < 0.05, n.s. not significant, Rank-sum test. **c**. Mean activation of all 25 cortical areas in naïve (gray) and expert (color) mice of L2/3 (black) and L5 (pink) during cue, pre, early-stim and late-stim periods. Areas are further divided into motor (orange), somatosensory + V1 (purple), association (green) and auditory (blue) areas. Error bars are s.e.m. across mice (n = 7 for each layer).

To highlight how activity patterns change with learning, we chose five cortical areas that captured the main learning-related modulations across layers (Fig. 2b). For each cortical area, we show a 2D response plot of trial vs. learning (Fig. 2bi; temporal profile of the trial on the x axis; learning profile in the y axis); trial responses during naïve and expert (Fig. 2bii); learning response during a specific trial period, what we term as a “learning curve” (Fig. 2biii); and an average response during each epoch, in naïve and expert animals (Fig. 2biv). These analyses show: first, that in both L2/3 and L5, BC displays similar and significant learning-related enhancement during the early-stim period (Fig. 2b left; p < 0.05; Signed-rank test). In contrast, activity in RL is significantly enhanced in L2/3 across learning but significantly decreases in L5 (Fig. 2b second from left; L5 shows significance slightly earlier in the cue-period; p < 0.05; Signed-rank test). A frame-by-frame statistical analysis is displayed in Supplementary Figure 2, which shows that suppression in L5 occurs slightly earlier than the enhancement in L2/3. The third area, S2, displays a similar laminar divergence to RL, with an emphasis on strong suppression specifically in L5 (Fig. 2b middle). In the fourth area, PL, we focused on the late-stim period, in which the mouse has already transformed the sensory information into motor output. During the late-stim period, PL in L5 is significantly increased across learning, whereas in L2/3 it is significantly decreased (Fig. 2b second from right; p < 0.05; Signed-rank test). The fifth area, M1, during the late-stim period, displays a significant increase across learning in L5 (p < 0.05; Signed-rank test) but not in L2/3 (p > 0.05; Signed-rank test; Fig. 2b right).

Next, we expanded our analysis to all 25 cortical areas and analyzed the mean response during each time period in naïve and expert phases for L2/3 and L5 (Fig. 2c). We highlight several learning-related modulations at a global scale: (1) A large-scale, learning-related suppression during the cue-period is evident in L5 (Figure 2c, Cue epoch). This effect is not observed in L2/3, which later shows enhanced activity in task-related areas during the pre-period, an effect not seen in L5. (2) During the late-stim period, L5 displays a widespread increase in expert animals, especially in the frontal cortex (i.e., M1, M2 etc.), whereas L2/3 activity, specifically in the posterior cortex, is suppressed and further decreases as mice become experts. These dynamics emphasize that as the mouse learns to transform sensory information into a motor response, L5 and L2/3 are dynamically and distinctly modulated.

To further investigate the interaction between learning dynamics and cortical activity, we defined a ‘Learning map’ for each time period (Figure 3). The learning curve (i.e., d′ as a function of trials) was correlated (using Pearson’s correlation, r) with the response profile (i.e., ΔF/F as a function of trials), for each pixel in the imaging area (Fig. 3a). A positive correlation value indicates that the response within the given pixel is correlated with learning, i.e., increases with learning. A negative correlation value indicates that the response within the given pixel decreases with learning. This analysis shows how the activity map in L2/3 and L5 changes progressively as mice learn. Example learning maps for one L2/3 and one L5 mouse show that during the cue-period, L5 is negatively correlated with learning across the whole cortex (Fig. 3b; Supplementary video 2). During the pre-period, L2/3 somatosensory and association area RL, are positively correlated with learning, whereas L5 activity across the majority of cortex remains negatively correlated. During the early-stim period, many sensory and motor areas including BC exhibit strong positive correlations with learning in both layers. During the late-stim period, L5 activity is positively correlated with learning across most of the cortex, whereas during this epoch, positive correlations for L2/3 are confined to the frontal cortex and negative correlations to the posterior cortex.

**Fig. 3.**
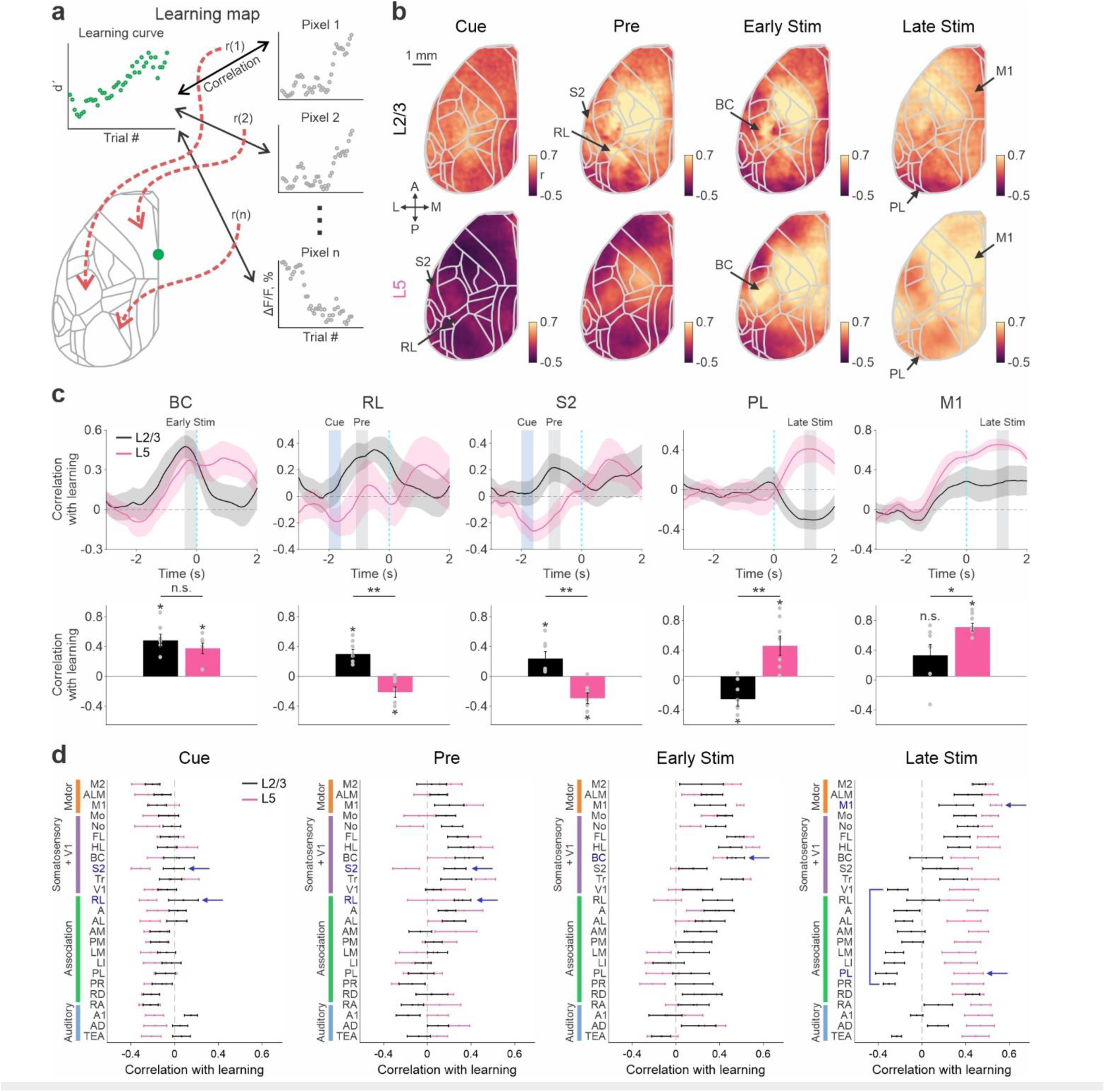
Learning maps reveal laminar temporal and activity dissociations in relationship to learning. **a** Schematic illustration for calculating a learning map. Each pixel in the map reflects the correlation coefficient (r) between the mouse’s learning curve and the response curve ΔF/F of that pixel across (termed as correlation with learning). This can be done within a specific time period (i.e., cue, pre, early-stim or late-stim) or for each time frame separately. **b** Learning maps during cue, pre, early-stim and late-stim periods in one example L2/3 (top) and L5 (bottom) mice. Color denotes r-values. Scale bar is 1 mm. **c** Top: correlation with learning as a function of time for the 5 selected areas (from left to right) averaged across mice for L2/3 (black; n = 7) and L5 (pink; n = 7). Error shading indicates ± s.e.m. Time period for analysis window is marked by gray or blue color. Bottom: Correlation with learning for each area during the specific trial period indicated at the top (blue bar is for L5). Error bars are s.e.m. across mice. Gray dots depict individual mice. *p < 0.05, ** p < 0.005, n.s. not significant, Rank-sum test. **d** Correlation with learning during cue, pre, early-stim and late-stim periods for all 25 cortical areas in L2/3 (black) and L5 (pink). Error bars are s.e.m. across mice.

Next, we examined the five representative cortical areas (as in Fig. 2b) and calculated their correlation with learning for each time frame separately (Fig. 3c). BC displayed significant positive correlations in both layers during the early-stim period (p < 0.05; Signed-rank test), with no significant difference between L2/3 and L5 (p > 0.05; Rank-sum test). In RL and S2 there is a divergence between layers: L5 is significantly negatively correlated with learning during the cue-period whereas L2/3 displays a significant positive correlation with learning at slightly later time points in the pre-period (Fig. 3c). In PL and M1, positive correlations are evident in L5 during the late-stim period. In this epoch, L2/3 in PL is significantly negatively correlated with learning, and M1 has no significant correlation with learning. A frame-by-frame statistical analysis between layers is presented in Supplementary Fig. 3.

Analysis across all 25 cortical areas (Fig. 3d), highlights the general negative correlation with learning in L5 during the cue-period. RL and S2 are two representative cortical areas that are differentially correlated, with a significant negative correlation in L5 during this epoch, and a significant positive correlation in L2/3 during the following pre-period. In the early-stim epoch, both L2/3 and L5 are similarly positively correlated with learning in somatosensory and motor areas. In the late-stim epoch, there is a significant positive correlation in L5, which is less pronounced in L2/3 anterior cortex and entirely absent in L2/3 posterior cortex, which displays negative correlations.

These results motivated us to develop a ‘Layer similarity index’ that could be used to quantify whether activity in each layer and each brain area changes similarly with learning. To do this, we defined a layer similarity index as the correlation of the learning traces with L2/3 and L5 in each cortical area were correlated to each other (Fig. 4a; values range from −1 to 1). A positive layer similarity index (r > 0) indicates that L2/3 and L5 display similar dynamics with regards to learning. A weak layer similarity index (r ∼ 0) indicates that L2/3 and L5 display different dynamics with regards to learning. A negative layer similarity index (r < 0) indicates that L2/3 and L5 display opposite dynamics with regards to learning. This analysis shows that the anterior sensory and motor areas: motor M2, ALM, and Somatosensory Mo, No, FL, HL, BC, and Trunk cortex changes similarly. Even some association areas (AM and RD) are modulated similarly during learning in L2/3 and L5 (further analyses to show the similar dynamics in M2 and RD between L2/3 and L5 are presented in Supplementary Fig. 4). In contrast, posterior lateral association areas (LM, LI, PL and PR) and V1, have negative layer similarity indices, i.e., opposing relations to learning for L2/3 and L5. M1, S2 and RL display a near zero index, i.e., different relations to learning for L2/3 and L5 (Fig. 4b; 4c). Taken together, these results suggest that L2/3 and L5 can have similar or different dynamics with regards to learning, depending on the cortical area.

**Fig. 4.**
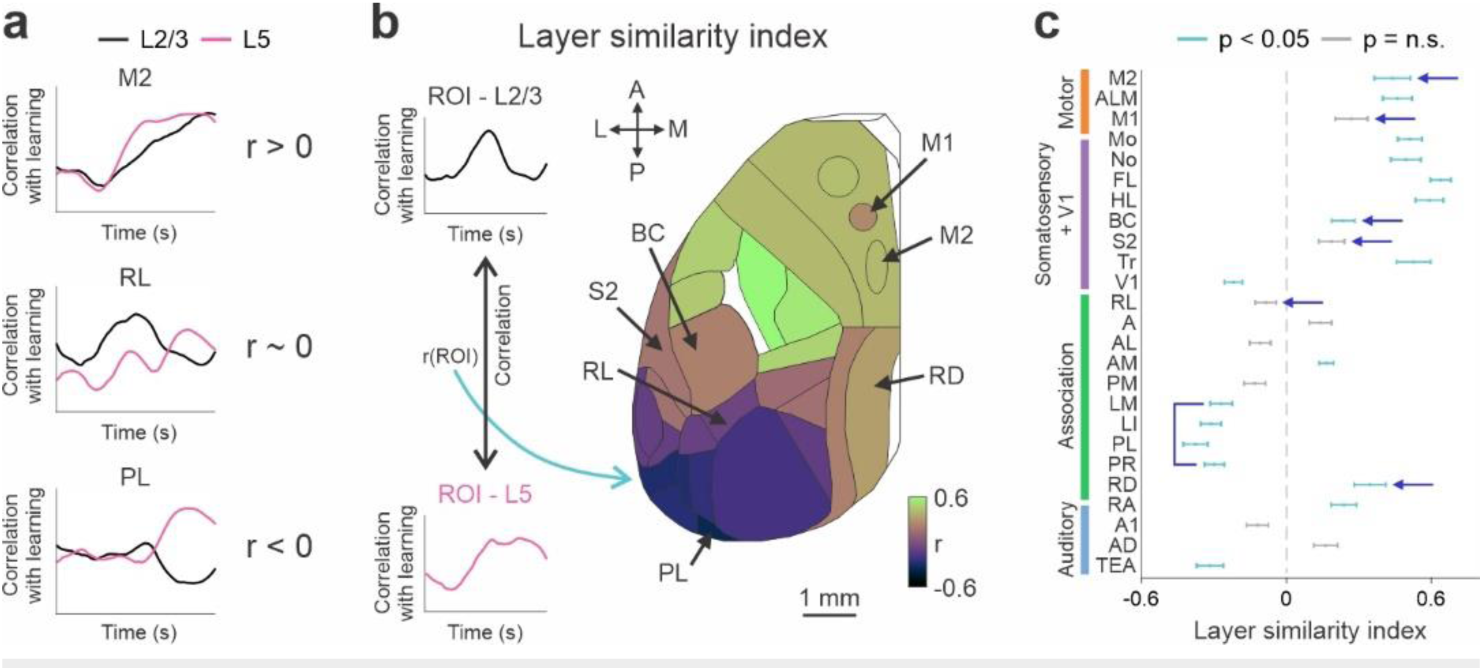
Layer similarity index reveals cortex-wide heterogeneity. **a** Example of three possible outcomes for the layer similarity index (r > 0, r ∼ 0 or r < 0) by correlating traces of correlation with learning (as in Fig. 3b) between L2/3 (black) and L5 (pink). **b** Left: Schematic for the calculation of layer similarity index. Right: Layer similarity index map. Positive values indicate similarity between L2/3 and L5 with regards to learning whereas negative values depict differences between layers. Scale bar is 1 mm. **c** Layer similarity index for all areas. Error bars are s.e.m. across mice. *p < 0.05 (green), n.s. not significant (gray), Signed-rank test.

### L5 temporal activity is strongly associated with task execution

Next, we analyzed the correlation between cortex-wide dynamics in L2/3 or L5 and movement. First, we defined a ‘Movement map’ by correlating the body movement trace with the response trace of each pixel in the imaged area (Fig. 5a). Data from each trial in naïve and expert cases for L2/3 and L5 were separately averaged. A positive value (r) indicates similar dynamics between the body movement and the response in a given pixel. A negative value indicates that body movement and the response in a given pixel had opposing dynamics. A movement map from a single animal reveal that in naïve L2/3 animals, there is a positive correlation between movement and pixel responses in frontal sensory and motor areas. This correlation for frontal areas becomes slightly stronger after learning (Fig. 5b). In naïve animals, the correlation to movement is weaker in posterior areas, and after learning, they are negatively correlated to movement. In L5, on the other hand, there is a positive correlation to movement throughout the cortex. This correlation further strengthens after learning, in the expert animal (Fig. 5b).

**Fig. 5.**
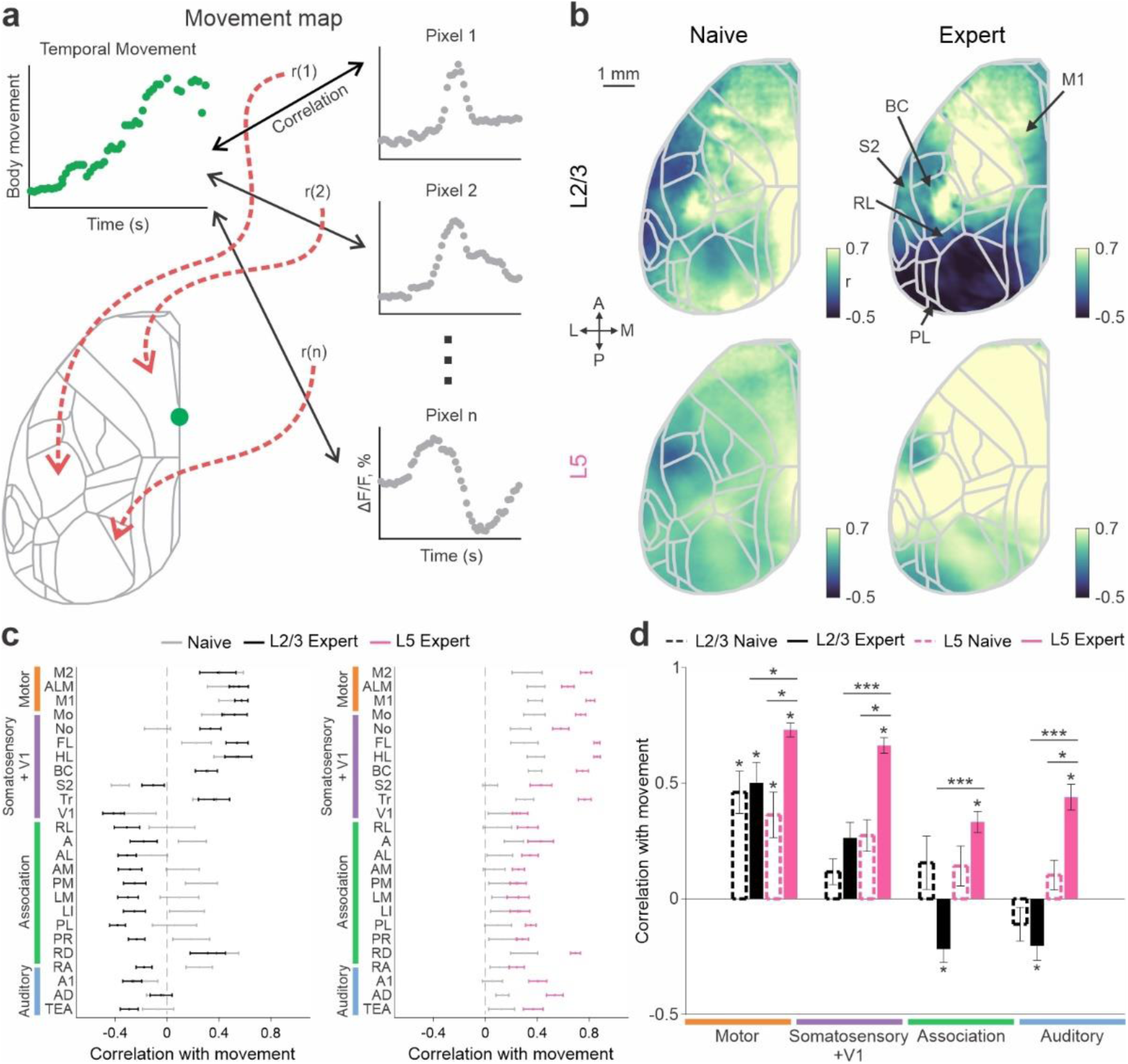
Correlation with movement reveals frontal/posterior divergence across layers. **a** Schematic illustration for calculating a Movement map. Each pixel in the maps reflects the correlation coefficient (r) between the mouse’s body movement and the temporal response (ΔF/F) of the respective pixel. This can be calculated for each trial and averaged across naïve or expert cases and for each layer separately. **b** Example movement maps in naïve (left) and expert (right) phase for L2/3 (top) and L5 (bottom). Color denotes r-values. Scale bar is 1 mm. **c** Correlation with movement in naïve (gray) and expert (color) for L2/3 (left; black) and L5 (right; pink) in all 25 areas. Error bars are s.e.m. across mice (n = 7 for each layer). **d** As in c, but grouped with motor (orange), somatosensory + V1 (purple), association (green) and auditory (blue) areas. *p < 0.05, *** p < 0.0005, Signed-rank test within layer, Rank-sum test across layers.

The average correlation maps between movement and activity across all 25 areas show a frontal/posterior dissociation for L2/3 (Fig. 5c). As the mice learn, the posterior cortex, excluding RD, becomes negatively correlated with movement. In L5, no such frontal/posterior divergence is observed. The correlation with movement is significantly higher in expert mice compared to naïve mice across all areas, with strong effect in the frontal cortex. This frontal/posterior divergence is also evident in the grouped data (Figure 5d; Motor, Somatosensory + V1,

Association and auditory cortex). Expert L5 mice display a positive correlation, particularly in the frontal cortex, whereas expert L2/3 mice exhibit a negative correlation with movement in posterior areas (Fig. 5d; p < 0.05; n=7 mice; Signed-rank test). Taken together, these results indicate that, with learning, L2/3 and L5 exhibit distinct cortex-wide dynamics in relation to body movement, in a lamina specific manner: L5 shows a positive movement correlation in frontal areas, while L2/3 shows a negative movement correlation in posterior cortex.

### Functional correlations between layers across temporal periods

Given this unique dataset of learning-related cortical dynamics across both layers, we aimed to assess the direct relationship between learning-related dynamics across layers. Specifically, we were interested in examining the link between the learning profile in one layer during a specific time period and the learning profile in another layer in a subsequent time period. To do this, we first defined a ‘ROI seed map’, in which we correlated the response learning curve (ΔF/F as a function of learning) in a cortical area, the seed area, with the response learning curve of each pixel across the 25 imaged areas (Fig. 6a; see also Methods). Positive correlation values indicate that, during learning, this pixel is modulated similarly to the seed area. Negative correlation values indicate that the pixel is modulated in the opposite direction to the seed area. This analysis can be performed for each layer individually, or by using an area in one layer (e.g., L5), it is possible to generate a map of activity in the other layer (e.g., L2/3). Additionally, this analysis can be performed for specific time epochs, for example, using the cue-period in the seed area and relating it to a different time period in the map, such as the subsequent pre-period (Fig. 6b).

**Fig. 6.**
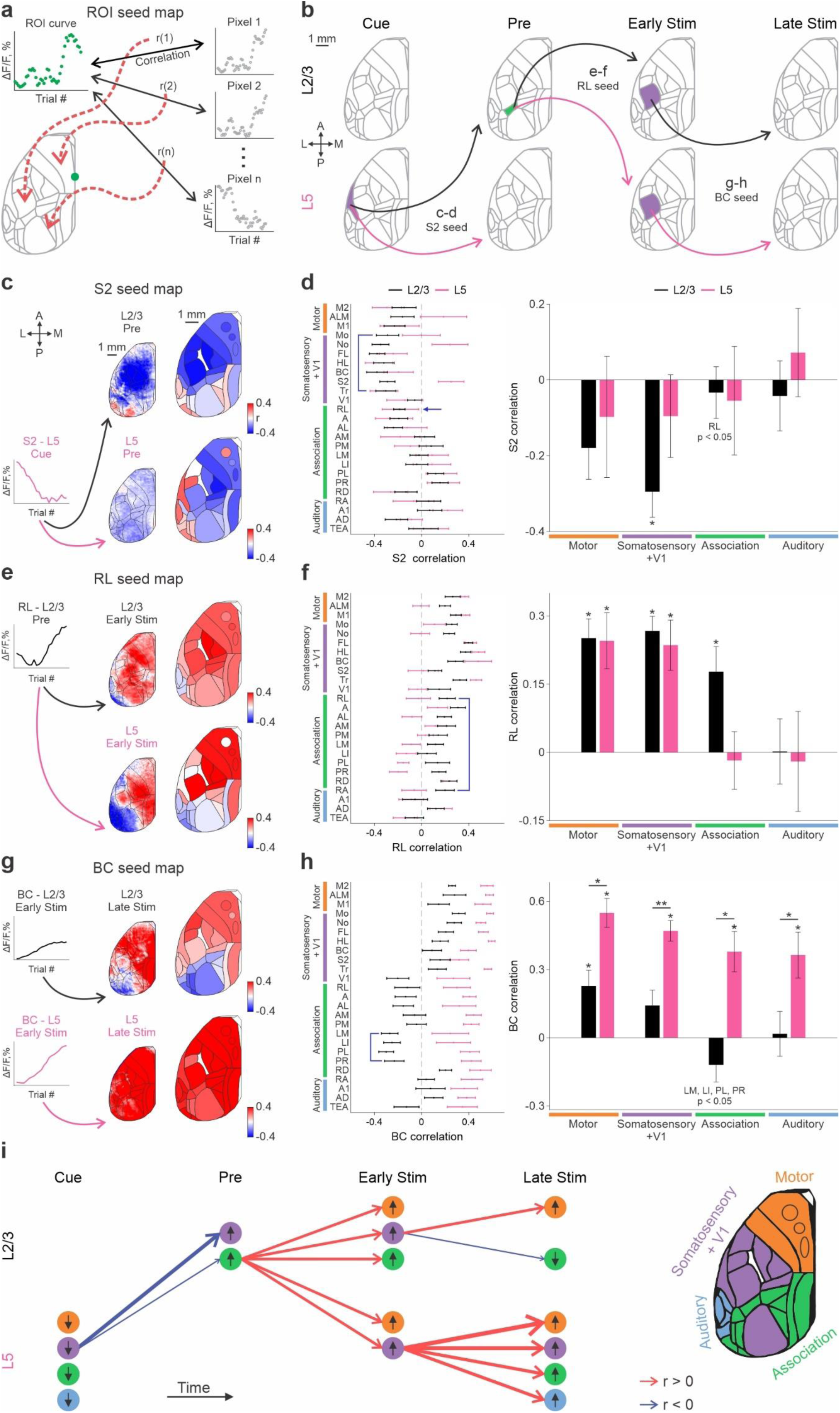
ROI seed maps between layers reveal cross-layer temporal dynamics. **a** Schematic for calculating a ROI seed map. Each pixel in the map reflects the correlation coefficient (r) between the learning-related response (ΔF/F) curve of a seed area (e.g. S2, RL, or BC) and the learning-related response (ΔF/F) curve of the respective pixel. This can be done within or across L2/3 and L5 and also across different time periods. **b** Schematic illustration of the ROI seed maps shown in panels c-h. A seed ROI in a certain layer and time period is correlated with the map of the subsequent time period in either L2/3 or L5. **c** ROI seed map between L5 S2 in the cue-period and the map of L2/3 (top) or L5 (bottom) in the subsequent pre-period. The small map is an example mouse and the big map is the average across mice (n = 7). **d** Left: Seed correlation values across all 25 cortical areas in L2/3 (black) and L5 (pink). Error bars depict mean ± s.e.m. across mice (n = 7). Right: Seed correlation values grouped into Motor, Somatosensory + V1, Association and Auditory for L2/3 and L5. **e,f** same as c,d but the seed is RL in L2/3 during the pre-period and the map is L2/3 or L5 during the early-stim period. **g,h** same as c,d but the seed area is BC either in L2/3 or L5 during the early-stim period and the map is either L2/3 or L5 during the late-stim period. *p < 0.05, **p < 0.005, Signed-rank test within layer, Rank-sum test across layers. **i** schematic summary of the above results. Each arrow depicts a correlation across time periods and layers where the amplitude is displayed as the width of the arrow and the sign is marked by color (blue or red for negative or positive correlations respectively). Each circle is a general region as in the inset on the right. Black arrows within each circle indicate the sign of response related to learning in that region.

Figure 6b illustrates specific relationships that are performed in this analysis. First, we calculated the relationship between the learning curve in S2 L5 during the cue-period and the activity map of L2/3 or L5 during the subsequent pre-period (Fig. 6c-d). This analysis reveals that during the cue-period, activity in S2 L5 is negatively correlated with most of the somatosensory cortex in L2/3 during the pre-period, and to a lesser extent, with L5 (Fig. 6c, d). Quantification across the 25 cortical areas and four areal groups in each layer highlights the negative values in L2/3 somatosensory areas (Fig. 6d; p<0.05; Signed-rank test). A seed analysis with each time frame instead of a specific epoch, as well as using S2 L2/3 as the seed area, is presented in Supplementary Figure 5a-c. These results suggest that as mice become experts, reduced activity in L5 of several cortical areas (including S2) during the cue-period leads to learning-related enhancement in L2/3 of task-related somatosensory areas (and RL).

To examine whether RL change in activity plays a role during the early-stim period as the texture approaches the whiskers, we repeated a similar analysis. We correlated the learning curve in RL L2/3 during the pre-period with activity maps of L2/3 or L5 during the subsequent early-stim period (Fig. 6e-f). During the pre-period, RL L2/3 activity is positively correlated with motor and somatosensory regions in both L2/3 and L5, and with association areas in L2/3 but not in L5 during the early-stim period (p < 0.05; Signed-rank test; Fig. 6f). A seed analysis with each time frame, as well as one using RL L5 as the seed area, is presented in Supplementary Figure 5d-f. These relationships between suggest that as the texture approaches the whiskers, a specific learning-related enhancement in RL opens a temporal window for subsequent sensory processing in somatosensory and motor cortex in both layers.

Next, we correlated the learning curve of BC during the early-stim period (either L2/3 or L5) with the activity maps of L2/3 or L5 during the late-stim period (Fig. 6g-h). BC L2/3 activity during the early-stim period is positively correlated with L2/3 motor areas and significantly negatively correlated in lateral posterior areas during late-stim period (PL, PR, LM and LI; p < 0.05; Signed-rank test). In contrast, BC L5 activity during the early-stim period is positively correlated with the entire dorsal L5 cortex during the late-stim period (p < 0.05; Signed-rank test). A frame-by-frame analysis highlights this divergence between layers, specifically during the late-stim period (Supplementary Fig. 5g). A complementary cross-layers analysis, switching between seed layers, reveals similar results (Supplementary Fig. 5h-i). These analyses suggest that BC increased activity during the early-stim period, integrating sensory information and transforming it into action. L5 population across the cortex is recruited during the late-stim period to execute the motor plan. Figure 6i summarizes our findings and highlights the main interlaminar interactions across the temporal profile of a Hit trial. In summary, we find cross-layer, learning-related interactions that may facilitate expert sensorimotor integration.

### L5 discriminates between choice prior to L2/3

Until now, we have focused on Hit trials throughout learning and their underlying cortex-wide dynamics in different layers. However, these analyses do not indicate whether activity in a certain cortical area discriminates between texture types. Does the discriminative power differ between L2/3 and L5? To address these questions, we computed the discrimination power between Hit and CR trials using receiver operating characteristic (ROC) analysis, with the area under the curve (AUC) representing discrimination power^1,44,48^ (Methods). AUC values range between 0 and 1, where extreme values of 0 or 1 indicate high levels of discriminative power. Here, we focused only on the expert case, due to the low probability in CR and Hit trials in the naïve case. In the current dataset, responses in Hit and CR trials during expert phase are relatively similar, with a tendency for higher responses in Hit trials emerging during the pre-period (Supplementary Fig. 6a-b). We applied this method to compute an AUC map for specific time periods, starting from the end of the cue-period to the end of the pre-period. The AUC maps from one example mouse of L2/3 and L5 reveal that the AUC first increases in BC and motor areas in L5 (−1.4 s, cue-pre) and only later becomes prominent in L2/3 (−1.0 s, pre-period; Fig.7a). We calculated the AUC for each time frame and grouped the AUC values into four regions. L5 exhibits a significant increase in AUC across all regions (Fig. 7b; p<0.05; Signed-rank test compared to 0.5). This change in AUC values preceded increases in L2/3, motor, somatosensory, and auditory areas. The peak of the AUC curve occurs earlier in L5 compared to L2/3 in motor and somatosensory areas (p<0.05; Rank-sum test). Averaging AUC values within specific time windows reveals significant AUC values (different than 0.5) in all regions between the cue and the pre periods (cue-pre) in L5 but not in L2/3 (Fig. 7c; p < 0.05; Signed-rank test). Later, during the pre-period, L2/3 joins L5 and develop significant AUC values in motor and somatosensory areas (p < 0.05; Signed-rank test). Taken together, both layers display discriminative power of choice (i.e., Hit vs. CR), with L5 slightly preceding L2/3 and exhibiting a more widespread discriminative capacity.

**Figure 7.**
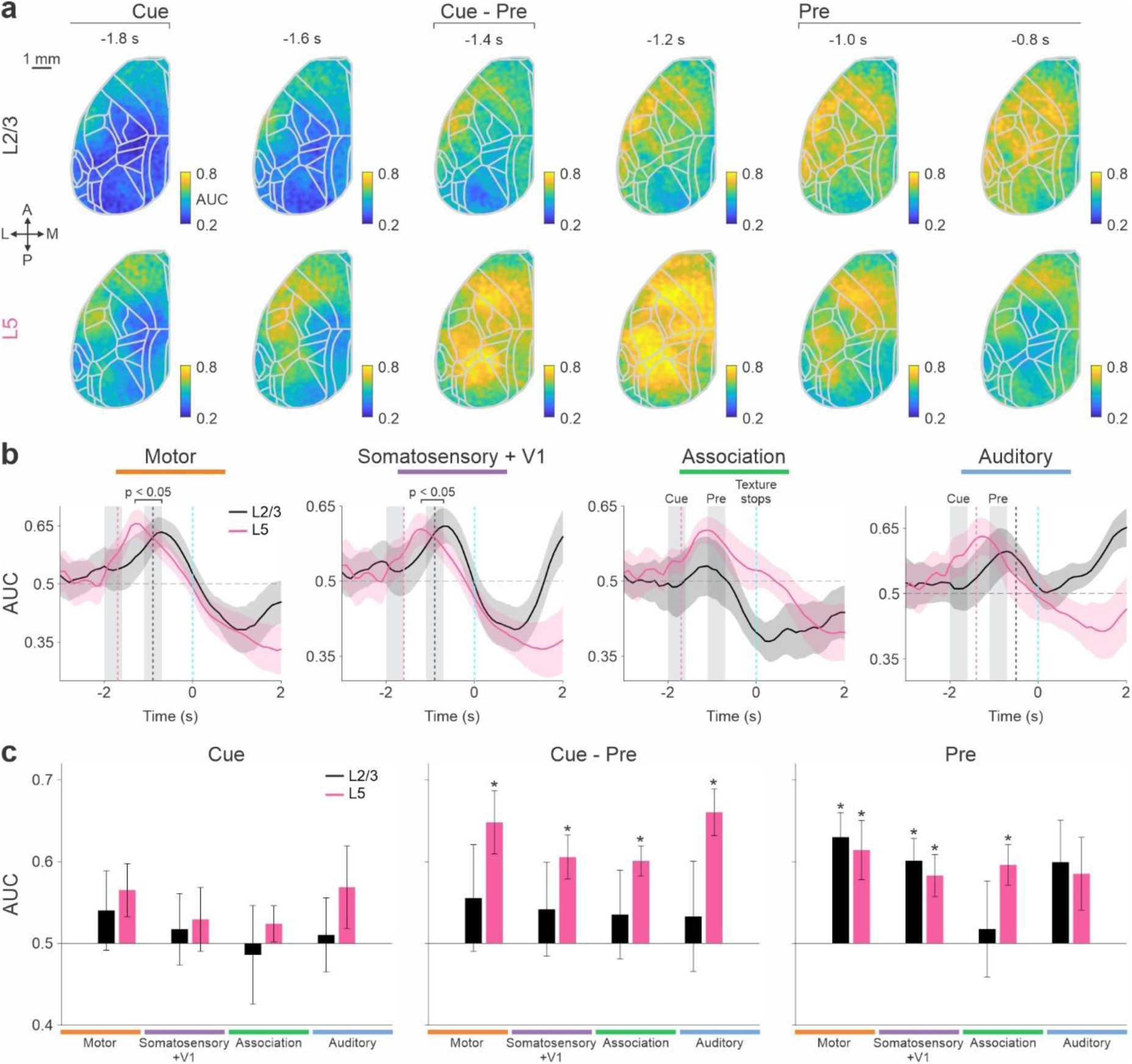
Discriminative power emerges in L5 prior to L2/3. **a** AUC maps (discriminating between Hit and CR trials) during cue to pre in one example expert mice of L2/3 (top) and L5 (bottom). Color denotes AUC values. Scale bar is 1 mm. **b** AUC values as a function of time for L2/3 (black) and L5 (pink) averaged across mice (n = 7). Dashed lines indicate first frame of AUC value with p < 0.05 (Signed-rank test) for L2/3 (black) and L5 (pink). Error shading indicates s.e.m. *p < 0.05, Rank-sum test of AUC peaks across layers. **c** AUC values during cue, cue-pre and pre in all areas grouped into motor (orange), somatosensory + V1 (purple), association (green) and auditory (blue) areas for L2/3 (black) and L5 (pink) mice. Error bars are s.e.m. across mice. *p < 0.05 Signed-rank test.

## Discussion

Here, we find that learning induces widespread, spatiotemporally complex changes in both L2/3 and L5. Surprisingly, learning-related dynamics were not always similar between layers and even displayed opposing dynamics in several higher-order cortical areas. Our main takeaway is that during learning, L5 plays a general role in suppressing cortex-wide responses before sensation and enhancing responses, particularly in frontal regions, after sensation during motor execution. In contrast, L2/3 contributes with local enhancement in task-related areas (in our case, RL and BC) just before and during sensation, thus enabling the optimal association of a certain stimulus with a reward.

### Laminar relationships within and across cortical areas

A widely accepted local organization is the cortical column which is considered a functional unit of information processing, having denser connections within the cortical column than between columns^50–53^. Whether different layers within a given cortical column change similarly or differently over days, as animals learn has been examined in very few contexts, especially for higher-order cortical areas. When observing the gradual modulation across the learning profile (i.e., across days), our results indicate inter-layer similarities or differences depending on the cortical area and the time along the trial (Fig. 4). For example, BC during the early sensation period and also M2 and RD, displayed similar learning-related dynamics in both L2/3 and L5. In contrast, several higher-order areas such as RL, S2 and PL display opposite learning dynamics across layers, in some cases L2/3 is enhanced and L5 is suppressed (e.g., RL and S2 during the pre-period) and vice versa in other cases (e.g., PL in late-stim period).

Our work suggests an update of the classical model of the canonical cortical circuit (L4 → L2/3 → L5/6)^54^ and the possible participation of other classes of local and global interactions. It is possible that L4-to-L5 inhibition, concurrent with L2/3 excitation during stimulus presentation sharpens sensory representations as mice learn and suppresses L5 activity^55–57^. It is also possible that direct thalamocortical input to deep layers is modified with learning independent of any changes in L2/3 activity^23,58–63^. L2/3 pyramidal neurons play a critical role in cortical processing by both directly exciting L5 pyramidal neurons and indirectly inhibiting them through activation of GABAergic neurons in L2/3 or L5^38–40^. Taken together, one or more of these circuit motifs might change gradually in the course of learning and modify activity in each cortical area in a unique fashion depending on task demands.

The exact balance between layers may be largely affected by diverse inputs from long-range cortical areas (e.g., S2, M1) subcortical areas (e.g., thalamus) and the neuromodulatory systems (e.g., cholinergic and adrenergic)^65–71^. Our work provides a global, cortex-wide window into activity in each pyramidal layer individually and their coactivation across cortical areas. We find strong extra-columnar interactions within and between L2/3 and L5 with a focus on subsequent time periods, e.g., from the cue to pre-period (Fig. 6c-d). As mice learn, activity in L5 of many cortical areas are negatively correlated with the learning profile of task-relevant somatosensory areas in L2/3 (during the pre-period). Another example, RL cortex in L2/3 (during pre-period) is positively correlated with the frontal cortex of L5 (during early-stim, but also in L2/3; Fig. 6e-f). Our results indicate that extra-columnar interactions can be stronger than within columnar interactions, emphasizing the importance of global laminar dynamics during learning. Indeed, several studies have highlighted the important roles of long-range projections from and onto specific layers, emphasizing top-down control^11,36,47,72–78^. For example, L2/3 in BC densely projects to L5A in motor cortex^36^, which may be recruited as the mouse learns to associate between the texture (i.e., sensory information encoded in BC) and the motor output (encoded in the frontal cortex). In summary, we find complex inter-laminar dynamics both locally (intra-columnar) and globally (extra-columnar) that may together promote learning.

### Global and local laminar dynamics underlying sensorimotor integration

In the current study mice learned to discriminate between two textures (sensed by their whiskers) and associate each one of them with a different outcome, reward or punishment. In rewarded trials, mice transformed the sensory information and association of each texture into a motor action (i.e., licking in Hit trials) in order to receive a reward. Furthermore, mice learned sensorimotor pattern is embedded within a fixed trial structure, starting with an auditory cue, followed by the texture approaching after which a texture moved into whisker touch and finally an auditory cue signaling the availability of reward (in response to a licking action). In a more general sense, mice learned to perform a sensorimotor integration within a fixed trial structure.

We find that both L2/3 and L5 population display similar sequential dynamics starting with a typical response in RL as the texture comes in, followed by BC activation during early sensation and ending with frontal cortex activity during motor execution (^1^; Fig. 2a). Nevertheless, learning-related modulations within the general trial structure and along the learning profile dramatically differ between L2/3 and L5. As the auditory is sounded, mice learn to link the cue with an incoming reward-related stimulus which requires sensory integration and discrimination (between go and no-go textures). In response to the cue, L5 populations decrease activity in a global manner with emphasis in RL, S2 and frontal cortex (Fig. 2C). In contrast, L2/3 displays minor learning-related suppression with local decreases in medial association areas^1^ (Fig. 2C). Although movement probability is only ∼20% (of all Hit trials) during the cue-period (Fig. 1h), mice (both L2/3 and L5) significantly decrease their movement after learning, indicating that the cue leads to quiescence in anticipation for the texture. Thus, global suppression in L5 may mediate a behavioral state of arousal in which motor planning and execution is diminished.

The laminar dynamics as a function of learning continue to change along the trial structure, as time goes by. As the texture comes into touch, learning-related modulation are transferred from global L5 suppression to local L2/3 enhancement, specifically in association area RL. RL enhancement in L2/3 just prior to sensation may convey task-related information such as attention and task history-related signals to Barrel cortex^18,79–81^. This top-down projection (specifically from L2/3) of higher-order information may make it possible for expert animals to optimally locate and discriminate between sensory stimuli. Sensory integration in BC occurs in both L2/3 and L5 and is likely to be related to a decision to lick or not, indicating that BC may be a key location in which sensation and action are associated (Fig. 2b). If the mouse decides to lick (i.e., execution), learning-related modulations are transformed from BC (presumably from L5) into a global signal in which L5 activity enhanced especially in the frontal cortex, whereas L2/3 is mostly suppressed in the posterior cortex. Taken together, these results highlight a sensorimotor pattern of activity that starts with global L5 suppression (in anticipation of an incoming sensation) followed by local sensory-related L2/3 enhancement (responsible for sensory integration) which is then transformed into global L5 enhancement (which represent a motor execution). This global (L5) – local (L2/3) – global (L5) sequence may be a fundamental neuronal process underlying sensorimotor integration.

### Frontal/posterior laminar divergence during movement execution

As discussed above, during late stages, L5 is globally enhanced with emphasis on the frontal cortex whereas L2/3 is suppressed in the posterior cortex. This divergence outlines a distinct frontal/posterior divergence across layers that emerges after learning (Fig. 4b). L5 frontal enhancement may represent the planning and execution of a motor action whereas L2/3 posterior suppression may indicate the active blocking of any additional incoming sensory information which may perturb the already ongoing sensorimotor integration. These results add on to the emerging evidence that frontal and posterior subnetworks are linked to different behavioral states, for example active and passive strategies respectively^32,48,82,83^. L2/3 suppression in the posterior cortex can be mediated by top-down feedback from frontal cortex^15,72,75,84,85^ (e.g., from M2) or subcortical areas (e.g., higher-order thalamus)^66,86^, targeting local inhibitory interneurons that effect primarily L2/3, e.g., VIP neurons^87^. Thus, as learning occurs, top-down inhibition may undergo plastic modulation and initiate as L5 frontal neurons become active during motor execution. Another mechanistic explanation for frontal/posterior divergence is the heterogenous effect of different neuromodulators, e.g., acetylcholine (ACh)^69,88^. ACh has been linked to aroused states and displays a frontal to posterior gradient across the dorsal cortex with optimal representation in the frontal cortex^88^. In addition, ACh differently affects cortical layers, primarily facilitating neuronal responses in L5 and suppressing L2/3^89–91^. Taken together, we propose that learning to transform a sensation into action promotes an ACh-mediated frontal motor cortex activation in L5 which projects back to posterior sensory cortex and suppresses L2/3 populations via local inhibitory circuits. These sequential dynamics (from frontal to posterior) could also explain the higher and earlier discrimination power in L5 compared to L2/3, implying that choice encoding may first emerge in deeper layers in the frontal cortex (^19^; Figure 7).

In summary, our work highlights how activity throughout cortex changes as animals learn a sensory motor task. This work reveals the intricate interactions that occur in each layer and each cortical area on every trial and over the course of days. We are just at the beginning of a profound understanding how cortical structure, function and interaction relate to the complex aspects of sensory motor integration and learning. Whether these changes in patterns of activity are causal for learning, whether and how these changes in patterns of activity relate to different classes of behavior and different classes of learning are still open to future investigations.

## Methods

### Animals

All experiments were approved by the Institutional Animal Care and Use Committee (IACUC) at the Hebrew University of Jerusalem, Israel (Permit Number: MD-20-16065-4). A total of 14 adult male mice (2-6 months old) were used in this study. Seven mice were triple-transgenic Rasgrf2-2A-dCre;CamK2a-tTA;TITL-GCaMP6f animals, expressing GCaMP6f in excitatory neocortical layer 2/3 neurons^1,90^ and seven were triple-transgenic Rbp4-dCre;CamK2a-tTA;TITL-GCaMP6f11, expressing GCaMP6f in excitatory neocortical layer 5 neurons, including both layer 5A and 5B neurons (Fig. 1c).

To generate L2/3 triple transgenic animals, double transgenic mice carrying CamK2a-tTa and TITL-GCaMP6f were crossed with a Rasgrf2-2A-dCre line. Individual lines are available from The Jackson Laboratory as JAX# 016198, JAX#024103, and JAX#022864, respectively). The Rasgrf2-2A-dCre;CamK2a-tTA;TITL-GCaMP6f line holds an inducible system in which the destabilized Cre (dCre) expressed under the control of the Rasgrf2-2A promoter, can be stabilized by trimethoprim (TMP) to be fully functional. TMP (Sigma T7883) was reconstituted in Dimethyl sulfoxide (DMSO, Sigma 34869) at a saturation level of 100 mg/ml, freshly prepared for each experiment. For TMP induction, mice were given a single intraperitoneal injection (150 µg TMP/g body weight; 29 g needle; 3–5 days post-surgery), diluted in 0.9% saline solution. To generate L5 triple transgenic animals, the Rbp4-dCre mouse line was crossed with the CamK2a-tTa;TITL-GCaMP6f double transgenic line. In this line, transgenic expression is suppressed by doxycycline treatment. These mice were not treated with doxycycline and therefore displayed strong expression throughout the experiment. Both lines display layer-specific expression homogeneously across the whole cortex with relatively minimal leakage into other layers, as previously reported (Fig. 1c;^90,91^).

### Surgery

We used an intact skull preparation for chronic wide-field calcium imaging of neocortical activity^1,26,45^. Mice were anesthetized with 2% isoflurane (in pure O_2_), and body temperature was maintained at 37 °C. We applied local analgesia (lidocaine 1%), exposed and cleaned the skull, and removed some muscles to access the entire dorsal surface of the left hemisphere (∼6 × 8 mm, from ∼3 mm anterior to bregma to ∼1 mm posterior to lambda; from the midline to at least 5 mm laterally). We built a wall around the hemisphere with adhesive material (G-Premio BOND; UV-cured) and dental cement “worms” (Charisma). Then, we applied transparent dental cement homogeneously over the imaging field (Tetric EvoFlow T1). Finally, a metal post for head fixation was glued to the back of the right hemisphere. This minimally invasive preparation enabled high-quality chronic imaging of neuronal population dynamics with a high success rate.

### Texture discrimination task

Mice were trained on a whisker-based go/no-go discrimination task (Fig. 1a) using a data acquisition interface (USB-6001; National Instruments) and custom-written LabVIEW software^1,18,92^. Each trial started with an auditory cue (stimulus cue; 2 beeps at 2 kHz, 100-ms duration with a 50-ms interval), signaling the approach of either one of two types of sandpapers (grit size P100: rough texture; P1200: smooth texture; 3M) to the mouse’s whiskers, as ‘go’ or ‘no-go’ textures (Fig. 1a). Sandpaper was mounted onto panels attached to a stepper motor (X-LMS100A; Zaber), which was mounted onto a motorized linear stage (T-LSM100A; Zaber) to move textures in and out of reach of the whiskers. The texture stayed in touch with the whiskers for 2 s, and then it was moved out after which an additional auditory cue (response cue; 4 beeps at 4 kHz, 50-ms duration with 25-ms interval) signaled the start of a 2 s response period. The stimulus and response cues were identical for both textures. A water reward (∼3 µL) was given to the mouse for licking for the go texture, but only after the response cue (‘Hit’), for the first correct lick during the response period (Fig. 1e; lick was detected using a piezo sensor). Punishment with white noise was given for licking in response to the no-go texture (‘false alarms’; FA). Licking before the response cue was neither rewarded nor punished. Rewards and punishments were omitted when mice withheld licking for the no-go (‘correct-rejections’, CR) or go (‘Miss’) textures. The licking detector remained in a fixed and reachable position throughout the entire trial. Licking before the response cue was allowed and did not lead to punishment or early reward.

### Training and performance

Five mice of each layer were trained to lick for the P100 texture (mice #1-4 and 7), and 2 mice were trained to lick for the P1200 texture (L2/3: mice #4 and 5, L5: mice #4 and 6). Mice were first handled and accustomed to head fixation before starting a water scheduling protocol. Before imaging began, mice were conditioned to lick for a reward after the go texture (presented within a similar trial structure as the task itself). Imaging began only after mice reliably licked following the response cue (typically after the first day; 200–400 trials). On the first day of imaging, mice were presented with the ‘go’ texture and after 50 trials the ‘no-go’ texture was gradually introduced (starting from 10% and increasing by 10% approximately every 50 trials^1^), until reaching 50% probability for the no-go texture by the end of the day. During the second day, most mice continuously licked for both textures (Supplementary Fig. 1a). After around 100 trials, we increased no-go probability to 80% and waited for mice to perform three continuous CR trials before returning to a 50% probability^93^. This was done for several rounds until mice increased their performance, specifically withheld licking for the no-go texture. In mice that continued to lick for both textures, we additionally repeated the wrong response until a correct response was made. In all mice, a 50% protocol was presented with no repetitions, as soon as they reached an expert level (d′ > 1.5). Overall, the training process took 5–8 consecutive days (ranging from 1223 to 2494 trials per mouse). An effort was made to maintain identical task parameters across days, e.g., the same texture position and location of the licking spout.

### Wide-field calcium imaging

The wide-field imaging setup is composed of a sensitive CMOS camera (Hamamatsu Orca Flash 4.0 v3; 512−512 pixels; 20 Hz) mounted on top of a dual-objective setup^26,47,94,95^. Two objectives (Navitar; D-5095, and D-2595) are interfaced with a dichroic (510 nm; AH) filter cube (Thorlabs). Blue LED light (Thorlabs; M470L3) is collimated and guided onto the preparation. Green light emitted from the preparation passes through both objectives before reaching the camera (at a 20 Hz frame rate). We also interleaved an additional 405 nm light used as a control for non-calcium related signals (by subtracting the normalized 405 signal from the normalized 473 signal; Thorlabs; M405L4; Teensy 4.0^18,25^). The frame rate was 20 Hz for the interleaved protocol, resulting in a 10 Hz corrected signal (see Data analysis below).

### Body movement monitoring and analysis

We used a body camera to detect general movements of the mouse (The Imaging Source DMK 33UX273; 30 Hz frame rate; Fig. 1g-h, Supplementary Fig. 1b). For each imaging day, we first outlined the forelimbs and the back areas (one area of interest for each), which were reliable areas for detecting general movements^1,18,26,47^. Next, we calculated the body movement (1 minus frame-to-frame correlation) within these areas as a function of time for each trial. We then averaged all the defined body areas into one ‘body’ vector. In general, we found that forelimb and back movement during task performance were highly correlated with excessive whisking of the mouse, which was validated by using an additional camera from the bottom to track whisker kinematics^18^.

### Data analysis

Data analysis was performed using Matlab software (Mathworks). All mice were continuously imaged during learning (5–8 days). Wide-field fluorescence images were downsampled down to 256×256 pixels, and pixels outside the imaging area were discarded. This resulted in a spatial resolution of ∼40 µm/pixel, which was sufficient to determine cortical borders in both L2/3 and L5, despite further scattering of emitted light through the tissue and skull. Using an interleaved imaging protocol, we first extracted signal to 470 nm and 405 nm traces. Next, for each trace, each pixel and each trial were normalized to the baseline from several frames before the stimulus cue (frame 0 division). Next, the 405 nm normalized traces were subtracted from the 473 nm normalized trace^18^. We report that this correction resulted in a minor modulation, mainly in reducing post-activation dips in the signal, but results were largely similar to those of mice imaged with only the 473 nm light^1,18,26^.

To define regions of interests (ROIs), the activation maps within each mouse were aligned together across days based on a semi-automatic protocol using blood vessel patterns to choose 5-7 points (cpselect function in MATLAB) and use them for transformation (cp2tform in MATLAB). Next, aligned activation maps were further registered onto the Allen atlas based on skull coordinates and functional patches (©2004 Allen Institute for Brain Science^1,46^, available from: http://mouse.brainmap.org/). Within the atlas borders, we defined 25 areas of interest, with some manual modifications within these borders to fit the functional activity for each mouse^1,18^. Motor cortex areas were defined based on stereotaxic coordinates and functional patches for each mouse (see below). Thus, all mice had similar regions of interest that were comparable within and across mice. Although the top 2D view for L5 may be slightly more condensed in lateral areas, we did not correct for this since we did not feel that L2/3 and L5 were substantially different (see Supplementary video 1). We further grouped these 25 areas into motor (orange), somatosensory + V1 (purple), association (green), and auditory (blue) areas (Fig. 1b). **Motor areas:** whisker-related primary motor cortex (M1; 1.5 anterior and 1 mm lateral from bregma, corresponding to the whisker evoked activation patch in M1 from the mapping session), anterior lateral motor cortex (ALM; 2.5 anterior and 1.5 mm lateral from bregma) and secondary motor cortex (M2); 1.5 anterior and 0.5 mm lateral from bregma corresponding. **Sensory areas:** somatosensory mouth (Mo), somatosensory nose (No), somatosensory hindlimb (HL), somatosensory forelimb (FL), barrel cortex (BC; primary somatosensory whisker); secondary somatosensory whisker (S2), somatosensory trunk (Tr), and primary visual cortex (V1). **Association cortex:** rostrolateral (RL), anterior (A), anterior lateral (AL), anterior medial (AM), posterior medial (PM), lateral medial (LM), lateral intermediate (LI), posterior lateral (PL), post-rhinal (PR), retrosplenial dorsal (RD), and retrosplenial angular (RA). **Auditory areas:** primary auditory (A1), auditory dorsal (AD), and temporal association area (TEA).

Within the temporal trial structure, we defined four time periods: cue (2 to 1.6 s before the texture stop) pre (1.1 to 0.7 s before the texture stop); early-stim (0.4 to 0 s before the texture stop); late-stim (1 to 1.4 s after the texture stop). Naïve and expert phases were defined in a window of 300 trials, approximately 150 trials before and after crossing the learning threshold.

The dataset consists of cortex-wide activation maps along the time course of a trial (7 seconds), repeated over the time course of learning (several days, hundreds of trials). This was collected for seven L2/3 mice and seven L5 mice. This three-dimensional dataset (brain x trial x learning^1^) can be presented in several ways: (1) Activity map (ΔF/F) averaged during a specific time period within a trial (i.e., cue, pre, early-stim or late-stim) and during expert or naïve cases (e.g., Fig 2a); (2) 2D plot of response in a certain cortical area where the x-axis is the trial structure and the y-axis is the learning profile (e.g., Fig 2bi); (3) A response curve across learning (i.e., a learning curve) during a specific time period within a trial for a certain brain area (Fig. 2biii); (4) A response during the trial averaged during naïve or expert cases for a certain brain area (e.g., Fig 2bii). Activity maps were smoothed with a Gaussian kernel (2σ = 7) for presentational purposes only.

### Calculation of behavioral learning curves

Trials were binned (n = 30 trials with no overlap) across learning, and the performance (defined as d′ = Z(Hit/(Hit + Miss)) – Z(FA/(FA + CR)) where Z denotes the inverse of the cumulative distribution function) was calculated for each bin. Next, each behavioral learning curve was fitted with a sigmoid function (sigmfit function in MATLAB^1^). A d′ = 1.5 was defined as the threshold for becoming expert (i.e. learning threshold; Fig. 1e).

### Learning maps and correlation with learning

To study the relationship between the behavioral learning curve and cortex-wide dynamics (either L2/3 or L5) across learning, we calculated a ‘Learning map’ (Fig. 3). This was done by calculating the Pearson’s correlation coefficient (r) between the behavioral learning curve of the mouse and the corresponding learning curve (i.e., response curve (ΔF/F) across learning) for each pixel (Fig. 3a). The r values were calculated for all pixels in the imaged area and can be presented as a map in which this is done for a specific time period (i.e. cue, pre, early-stim or late-stim period; Fig. 3b; Supplementary video 2) or for each time frame (Fig. 3c). Positive r values depict similar dynamics in a given brain area to the behavioral performance of the mouse. Negative r values depict opposing dynamics in a given brain area to the behavioral performance of the mouse (i.e., a decrease in activity along the learning profile). To quantify similarities and differences between L2/3 and L5 with relation to learning, we defined a ‘Layer similarity index’ (Fig. 4). The layer similarity index was defined as the Pearson’s correlation coefficient (r) between L2/3 and L5 correlations with learning traces (Fig. 3c and Fig. 4a, b). A positive layer similarity index (r > 0) indicates that L2/3 and L5 display similar dynamics with regard to learning. A negative layer similarity index (r < 0) indicates that L2/3 and L5 display different dynamics with regard to learning. The above analyses were done for each mouse separately and then averaged across either L2/3 or L5 mice.

### Movement maps and correlation with movement

To study the relationship between the body movement of mice and the accompanying cortex-wide dynamics in L2/3 and L5, we calculated a ‘Movement map’. This was done by calculating the Pearson’s correlation coefficient (r) between the behavioral movement vector (along the trial) of the mouse and the neuronal response (along the trial) in each pixel (Fig. 5a). This was done for each trial and then can be averaged within the naïve and expert phases (Fig. 5b). Positive r values depict similar dynamics between the body movement and the response in a given pixel along the trial. Negative r values depict opposing dynamics between the body movement and the response in a given pixel along the trial. The r values can then be averaged into 25 areas (Fig. 5c) or into four general regions (Fig. 5d).

### Regions of interests seed maps across layers and across time periods

To study the relationship between the learning curves of different layers and different time periods, we calculated a ‘ROI seed map’. In general, a ROI seed map calculates the Pearson’s correlation coefficient (r) between the learning curve (i.e., the response across the learning profile) of a certain area (i.e., seed region of interest) and the learning curve of each pixel in the imaged area (Fig. 6a). Positive r values indicate similar learning-related response dynamics between the seed areas and the given pixel. Negative r values indicate opposing learning-related response dynamics between the seed areas and the given pixel. Next, ROI seed maps can be calculated across different layers, for example the seed learning curve from L5 with the map (i.e., pixels) from L2/3. In addition, a ROI seed map can be taken from different time periods within the trial, for example, the seed learning curve can be during the cue-period and the map can be from the pre-period. Figure 6b displays the different types of ROI seed maps that are presented in this study, with emphasis on studying cross-layer interactions along subsequent time periods. To achieve a statistical comparison between layers, the learning curve of a seed ROI in a certain layer was correlated with the map of each mouse in the other layer, thus achieving n = 7 mice, which was used for statistical comparison.

### Discrimination power between Hit and CR trials in different layers

To measure how well neuronal populations could discriminate between Hit and CR trials during the expert phase, we calculated a receiver operating characteristic (ROC) curve between the response in Hit trials and CR trials (ΔF/F) and calculated its area under the curve (AUC) to determine the accuracy of the classifier. AUC values range from 0 to 1, where values around 0.5 indicate no discrimination power and extreme values (near 1 or 0) indicate high discriminative power. This analysis was done for each pixel in the imaging area (averaged during a specific time period), thus, obtaining an AUC map (Fig. 7a). In addition, AUC can be calculated for each time frame separately (Fig. 7b) and averaged for each cortical area or general region (Fig. 7c).

## Statistical analysis

In general, non-parametric two-tailed statistical tests were used: Rank-sum test to compare the medians of two populations, or the Signed-rank test to compare a population’s median to zero (or between two paired populations). Our statistical comparisons were performed at the mouse level with a sample of n = 7 for each layer.

## Author contributions

Y.E.P and A.G conceptualized the study. Y.E.P performed the experiments, preprocessed and analyzed the data. Y.E.P and A.G wrote the manuscript, and all authors revised, read and approved the final version of the manuscript.

## Data availability

Data will be shared upon reasonable request by the corresponding author. No new material was generated in this work.

## Code availability

Custom codes for data analysis were written in MATLAB and are available from the corresponding author upon request.

## Supplementary materials

Fig. S1 to S6. Vid. V1 and V2.

## Acknowledgements

We would like to thank Malak Abumadi and Dr. Odeya Marmor for their help with genotyping and mouse maintenance, Fritjof Helmchen and Philipp Bethge for the help with the transgenic mouse lines. This work is funded by the Einstein Foundation Research Grant (A-2021-644; A.G and M.L) and the European Union (ERC Starting Grant, MESO-AG, 101040378).

## References

1. Gilad, A., and Helmchen, F. (2020). Spatiotemporal refinement of signal flow through association cortex during learning. Nat Commun 11, 1744. 10.1038/s41467-020-15534-z.

2. Blake, D.T., Strata, F., Churchland, A.K., and Merzenich, M.M. (2002). Neural correlates of instrumental learning in primary auditory cortex. Proceedings of the National Academy of Sciences 99, 10114–10119. 10.1073/pnas.092278099.

3. Poort, J., Khan, A.G., Pachitariu, M., Nemri, A., Orsolic, I., Krupic, J., Bauza, M., Sahani, M., Keller, G.B., Mrsic-Flogel, T.D., et al. (2015). Learning Enhances Sensory and Multiple Non-sensory Representations in Primary Visual Cortex. Neuron 86, 1478–1490. 10.1016/j.neuron.2015.05.037.

4. Meister, M. (2022). Learning, fast and slow. Curr Opin Neurobiol 75, 102555. 10.1016/j.conb.2022.102555.

5. Carandini, M., and Churchland, A.K. (2013). Probing perceptual decisions in rodents. Nat Neurosci 16, 824–831. 10.1038/nn.3410.

6. Drieu, C., Zhu, Z., Wang, Z., Fuller, K., Wang, A., Elnozahy, S., and Kuchibhotla, K. (2024). Rapid emergence of latent knowledge in the sensory cortex drives learning. Preprint, 10.1101/2024.06.10.597946.

7. Pho, G.N., Goard, M.J., Woodson, J., Crawford, B., and Sur, M. (2018). Task-dependent representations of stimulus and choice in mouse parietal cortex. Nat Commun 9, 2596. 10.1038/s41467-018-05012-y.

8. Esmaeili, V., Tamura, K., Muscinelli, S.P., Modirshanechi, A., Boscaglia, M., Lee, A.B., Oryshchuk, A., Foustoukos, G., Liu, Y., Crochet, S., et al. (2021). Rapid suppression and sustained activation of distinct cortical regions for a delayed sensory-triggered motor response. Neuron 109, 2183–2201.e9. 10.1016/j.neuron.2021.05.005.

9. Roelfsema, P.R., and Holtmaat, A. (2018). Control of synaptic plasticity in deep cortical networks. Nat Rev Neurosci 19, 166–180. 10.1038/nrn.2018.6.

10. Ohl, F.W., and Scheich, H. (2005). Learning-induced plasticity in animal and human auditory cortex. Curr Opin Neurobiol 15, 470–477. 10.1016/j.conb.2005.07.002.

11. Chen, J.L., Margolis, D.J., Stankov, A., Sumanovski, L.T., Schneider, B.L., and Helmchen, F. (2015). Pathway-specific reorganization of projection neurons in somatosensory cortex during learning. Nat Neurosci 18, 1101–1108. 10.1038/nn.4046.

12. Li, W., Piëch, V., and Gilbert, C.D. (2008). Learning to Link Visual Contours. Neuron 57, 442–451. 10.1016/j.neuron.2007.12.011.

13. Han, S., and Helmchen, F. (2024). Coordinated multi-level adaptations across neocortical areas during task learning. Preprint, 10.1101/2024.09.26.615162.

14. Chia, X.W., Tan, J.K., Ang, L.F., Kamigaki, T., and Makino, H. (2023). Emergence of cortical network motifs for short-term memory during learning. Nat Commun 14, 6869. 10.1038/s41467-023-42609-4.

15. Makino, H., and Komiyama, T. (2015). Learning enhances the relative impact of top-down processing in the visual cortex. Nat Neurosci 18, 1116–1122. 10.1038/nn.4061.

16. Esmaeili, V., Oryshchuk, A., Asri, R., Tamura, K., Foustoukos, G., Liu, Y., Guiet, R., Crochet, S., and Petersen, C.C.H. (2022). Learning-related congruent and incongruent changes of excitation and inhibition in distinct cortical areas. PLoS Biol 20, e3001667. 10.1371/journal.pbio.3001667.

17. Makino, H., Ren, C., Liu, H., Kim, A.N., Kondapaneni, N., Liu, X., Kuzum, D., and Komiyama, T. (2017). Transformation of Cortex-wide Emergent Properties during Motor Learning. Neuron 94, 880–890.e8. 10.1016/j.neuron.2017.04.015.

18. Marmor, O., Pollak, Y., Doron, C., Helmchen, F., and Gilad, A. (2023). History information emerges in the cortex during learning. Elife 12. 10.7554/eLife.83702.

19. Steinfeld, R., Tacão-Monteiro, A., and Renart, A. (2024). Differential representation of sensory information and behavioral choice across layers of the mouse auditory cortex. Current Biology 34, 2200–2211.e6. 10.1016/j.cub.2024.04.040.

20. Naka, A., and Adesnik, H. (2016). Inhibitory Circuits in Cortical Layer 5. Front Neural Circuits 10. 10.3389/fncir.2016.00035.

21. Ren, C., Peng, K., Yang, R., Liu, W., Liu, C., and Komiyama, T. (2022). Global and subtype-specific modulation of cortical inhibitory neurons regulated by acetylcholine during motor learning. Neuron 110, 2334–2350.e8. 10.1016/j.neuron.2022.04.031.

22. Khan, A.G., Poort, J., Chadwick, A., Blot, A., Sahani, M., Mrsic-Flogel, T.D., and Hofer, S.B. (2018). Distinct learning-induced changes in stimulus selectivity and interactions of GABAergic interneuron classes in visual cortex. Nat Neurosci 21, 851–859. 10.1038/s41593-018-0143-z.

23. Constantinople, C.M., and Bruno, R.M. (2013). Deep Cortical Layers Are Activated Directly by Thalamus. Science (1979) 340, 1591–1594. 10.1126/science.1236425.

24. Economo, M.N., Viswanathan, S., Tasic, B., Bas, E., Winnubst, J., Menon, V., Graybuck, L.T., Nguyen, T.N., Smith, K.A., Yao, Z., et al. (2018). Distinct descending motor cortex pathways and their roles in movement. Nature 563, 79–84. 10.1038/s41586-018-0642-9.

25. Musall, S., Kaufman, M.T., Juavinett, A.L., Gluf, S., and Churchland, A.K. (2019). Single-trial neural dynamics are dominated by richly varied movements. Nat Neurosci 22, 1677–1686. 10.1038/s41593-019-0502-4.

26. Gilad, A., Gallero-Salas, Y., Groos, D., and Helmchen, F. (2018). Behavioral Strategy Determines Frontal or Posterior Location of Short-Term Memory in Neocortex. Neuron 99, 814–828.e7. 10.1016/j.neuron.2018.07.029.

27. Stringer, C., Pachitariu, M., Steinmetz, N., Reddy, C.B., Carandini, M., and Harris, K.D. (2019). Spontaneous behaviors drive multidimensional, brainwide activity. Science (1979) 364. 10.1126/science.aav7893.

28. Musall, S., Sun, X.R., Mohan, H., An, X., Gluf, S., Li, S.-J., Drewes, R., Cravo, E., Lenzi, I., Yin, C., et al. (2023). Pyramidal cell types drive functionally distinct cortical activity patterns during decision-making. Nat Neurosci. 10.1038/s41593-022-01245-9.

29. Cheung, J.A., Maire, P., Kim, J., Lee, K., Flynn, G., and Hires, S.A. (2020). Independent representations of self-motion and object location in barrel cortex output. PLoS Biol 18, e3000882. 10.1371/journal.pbio.3000882.

30. Hubel, D.H., and Wiesel, T.N. (1963). Shape and arrangement of columns in cat’s striate cortex. J Physiol 165, 559–568. 10.1113/jphysiol.1963.sp007079.

31. Mountcastle, V.B., Davies, P.W., and Berman, A.L. (1957). RESPONSE PROPERTIES OF NEURONS OF CAT’S SOMATIC SENSORY CORTEX TO PERIPHERAL STIMULI. J Neurophysiol 20, 374–407. 10.1152/jn.1957.20.4.374.

32. Mao, T., Kusefoglu, D., Hooks, B.M., Huber, D., Petreanu, L., and Svoboda, K. (2011). Long-Range Neuronal Circuits Underlying the Interaction between Sensory and Motor Cortex. Neuron 72, 111–123. 10.1016/j.neuron.2011.07.029.

33. Onodera, K., and Kato, H.K. (2022). Translaminar recurrence from layer 5 suppresses superficial cortical layers. Nat Commun 13, 2585. 10.1038/s41467-022-30349-w.

34. Lefort, S., Tomm, C., Floyd Sarria, J.-C., and Petersen, C.C.H. (2009). The Excitatory Neuronal Network of the C2 Barrel Column in Mouse Primary Somatosensory Cortex. Neuron 61, 301–316. 10.1016/j.neuron.2008.12.020.

35. Petreanu, L., Huber, D., Sobczyk, A., and Svoboda, K. (2007). Channelrhodopsin-2–assisted circuit mapping of long-range callosal projections. Nat Neurosci 10, 663–668. 10.1038/nn1891.

36. Adesnik, H., and Scanziani, M. (2010). Lateral competition for cortical space by layer-specific horizontal circuits. Nature 464, 1155–1160. 10.1038/nature08935.

37. HUBEL, D.H., and WIESEL, T.N. (2004). Brain and Visual Perception (Oxford University PressNew York) 10.1093/acprof:oso/9780195176186.001.0001.

38. Harris, K.D., and Shepherd, G.M.G. (2015). The neocortical circuit: themes and variations. Nat Neurosci 18, 170–181. 10.1038/nn.3917.

39. Hwang, E.J., Dahlen, J.E., Mukundan, M., and Komiyama, T. (2017). History-based action selection bias in posterior parietal cortex. Nat Commun 8, 1242. 10.1038/s41467-017-01356-z.

40. Masamizu, Y., Tanaka, Y.R., Tanaka, Y.H., Hira, R., Ohkubo, F., Kitamura, K., Isomura, Y., Okada, T., and Matsuzaki, M. (2014). Two distinct layer-specific dynamics of cortical ensembles during learning of a motor task. Nat Neurosci 17, 987–994. 10.1038/nn.3739.

41. Moberg, S., Garibbo, M., Mazo, C., Gilad, A., Schmitz, D., Costa, R.P., Larkum, M.E., and Takahashi, N. (2025). Distinct roles of cortical layer 5 subtypes in associative learning. Preprint, 10.1101/2025.01.07.631500.

42. Lacefield, C.O., Pnevmatikakis, E.A., Paninski, L., and Bruno, R.M. (2019). Reinforcement Learning Recruits Somata and Apical Dendrites across Layers of Primary Sensory Cortex. Cell Rep 26, 2000–2008.e2. 10.1016/j.celrep.2019.01.093.

43. Benezra, S.E., Patel, K.B., Perez Campos, C., Hillman, E.M., and Bruno, R.M. (2024). Learning enhances behaviorally relevant representations in apical dendrites. Elife 13. 10.7554/eLife.98349.

44. Chen, J.L., Carta, S., Soldado-Magraner, J., Schneider, B.L., and Helmchen, F. (2013). Behaviour-dependent recruitment of long-range projection neurons in somatosensory cortex. Nature 499, 336–340. 10.1038/nature12236.

45. Silasi, G., Xiao, D., Vanni, M.P., Chen, A.C.N., and Murphy, T.H. (2016). Intact skull chronic windows for mesoscopic wide-field imaging in awake mice. J Neurosci Methods 267, 141–149. 10.1016/j.jneumeth.2016.04.012.

46. Oh, S.W., Harris, J.A., Ng, L., Winslow, B., Cain, N., Mihalas, S., Wang, Q., Lau, C., Kuan, L., Henry, A.M., et al. (2014). A mesoscale connectome of the mouse brain. Nature 508, 207–214. 10.1038/nature13186.

47. Gallero-Salas, Y., Han, S., Sych, Y., Voigt, F.F., Laurenczy, B., Gilad, A., and Helmchen, F. (2021). Sensory and Behavioral Components of Neocortical Signal Flow in Discrimination Tasks with Short-Term Memory. Neuron 109, 135–148.e6. 10.1016/j.neuron.2020.10.017.

48. Gilad, A., Meirovithz, E., and Slovin, H. (2013). Population Responses to Contour Integration: Early Encoding of Discrete Elements and Late Perceptual Grouping. Neuron 78, 389–402. 10.1016/j.neuron.2013.02.013.

49. Mountcastle, V.B. (1957). MODALITY AND TOPOGRAPHIC PROPERTIES OF SINGLE NEURONS OF CAT’S SOMATIC SENSORY CORTEX. J Neurophysiol 20, 408–434. 10.1152/jn.1957.20.4.408.

50. Mountcastle, V. (1997). The columnar organization of the neocortex. Brain 120, 701–722. 10.1093/brain/120.4.701.

51. Bennett, M. (2020). An Attempt at a Unified Theory of the Neocortical Microcircuit in Sensory Cortex. Front Neural Circuits 14. 10.3389/fncir.2020.00040.

52. Buxhoeveden, D.P., and Casanova, M.F. (2002). The minicolumn hypothesis in neuroscience. Brain 125, 935–951. 10.1093/brain/awf110.

53. Douglas, R.J., and Martin, K.A.C. (2004). NEURONAL CIRCUITS OF THE NEOCORTEX. Annu Rev Neurosci 27, 419–451. 10.1146/annurev.neuro.27.070203.144152.

54. Onodera, K., and Kato, H.K. (2022). Translaminar recurrence from layer 5 suppresses superficial cortical layers. Nat Commun 13, 2585. 10.1038/s41467-022-30349-w.

55. Naka, A., and Adesnik, H. (2016). Inhibitory Circuits in Cortical Layer 5. Front Neural Circuits 10. 10.3389/fncir.2016.00035.

56. Pluta, S., Naka, A., Veit, J., Telian, G., Yao, L., Hakim, R., Taylor, D., and Adesnik, H. (2015). A direct translaminar inhibitory circuit tunes cortical output. Nat Neurosci 18, 1631–1640. 10.1038/nn.4123.

57. Bureau, I., von Saint Paul, F., and Svoboda, K. (2006). Interdigitated Paralemniscal and Lemniscal Pathways in the Mouse Barrel Cortex. PLoS Biol 4, e382. 10.1371/journal.pbio.0040382.

58. Freund, T.F., Martin, K.A.C., and Whitteridge, D. (1985). Innervation of cat visual areas 17 and 18 by physiologically identified X- and Y-type thalamic afferents. I. Arborization patterns and quantitative distribution of postsynaptic elements. Journal of Comparative Neurology 242, 263–274. 10.1002/cne.902420208.

59. Humphrey, A.L., Sur, M., Uhlrich, D.J., and Sherman, S.M. (1985). Termination patterns of individual X- and Y-cell axons in the visual cortex of the cat: Projections to area 18, to the 17/18 border region, and to both areas 17 and 18. Journal of Comparative Neurology 233, 190–212. 10.1002/cne.902330204.

60. Wimmer, V.C., Bruno, R.M., de Kock, C.P.J., Kuner, T., and Sakmann, B. (2010). Dimensions of a Projection Column and Architecture of VPM and POm Axons in Rat Vibrissal Cortex. Cerebral Cortex 20, 2265–2276. 10.1093/cercor/bhq068.

61. Huang, W., Armstrong-James, M., Rema, V., Diamond, M.E., and Ebner, F.F. (1998). Contribution of Supragranular Layers to Sensory Processing and Plasticity in Adult Rat Barrel Cortex. J Neurophysiol 80, 3261–3271. 10.1152/jn.1998.80.6.3261.

62. Schwark, H.D., Malpeli, J.G., Weyand, T.G., and Lee, C. (1986). Cat area 17. II. Response properties of infragranular layer neurons in the absence of supragranular layer activity. J Neurophysiol 56, 1074–1087. 10.1152/jn.1986.56.4.1074.

63. Pouille, F., and Scanziani, M. (2004). Routing of spike series by dynamic circuits in the hippocampus. Nature 429, 717–723. 10.1038/nature02615.

64. Neske, G.T., and Cardin, J.A. (2025). Higher-order thalamic input to cortex selectively conveys state information. Cell Rep 44, 115292. 10.1016/j.celrep.2025.115292.

65. Pardi, M.B., Vogenstahl, J., Dalmay, T., Spanò, T., Pu, D.-L., Naumann, L.B., Kretschmer, F., Sprekeler, H., and Letzkus, J.J. (2020). A thalamocortical top-down circuit for associative memory. Science (1979) 370, 844–848. 10.1126/science.abc2399.

66. Cruz, K.G., Leow, Y.N., Le, N.M., Adam, E., Huda, R., and Sur, M. (2023). Cortical-subcortical interactions in goal-directed behavior. Physiol Rev 103, 347–389. 10.1152/physrev.00048.2021.

67. Slater, B.J., Sons, S.K., Yudintsev, G., Lee, C.M., and Llano, D.A. (2019). Thalamocortical and Intracortical Inputs Differentiate Layer-Specific Mouse Auditory Corticocollicular Neurons. The Journal of Neuroscience 39, 256–270. 10.1523/JNEUROSCI.3352-17.2018.

68. Collins, L., Francis, J., Emanuel, B., and McCormick, D.A. (2023). Cholinergic and noradrenergic axonal activity contains a behavioral-state signal that is coordinated across the dorsal cortex. Elife 12. 10.7554/eLife.81826.

69. Yang, H., Kwon, S.E., Severson, K.S., and O’Connor, D.H. (2016). Origins of choice-related activity in mouse somatosensory cortex. Nat Neurosci 19, 127–134. 10.1038/nn.4183.

70. Ni, J., and Chen, J.L. (2017). Long-range cortical dynamics: a perspective from the mouse sensorimotor whisker system. European Journal of Neuroscience 46, 2315–2324. 10.1111/ejn.13698.

71. Zhang, S., Xu, M., Chang, W.-C., Ma, C., Hoang Do, J.P., Jeong, D., Lei, T., Fan, J.L., and Dan, Y. (2016). Organization of long-range inputs and outputs of frontal cortex for top-down control. Nat Neurosci 19, 1733–1742. 10.1038/nn.4417.

72. Zhang, S., Xu, M., Kamigaki, T., Hoang Do, J.P., Chang, W.-C., Jenvay, S., Miyamichi, K., Luo, L., and Dan, Y. (2014). Long-range and local circuits for top-down modulation of visual cortex processing. Science (1979) 345, 660–665. 10.1126/science.1254126.

73. D’Souza, R.D., and Burkhalter, A. (2017). A Laminar Organization for Selective Cortico-Cortical Communication. Front Neuroanat 11. 10.3389/fnana.2017.00071.

74. Schneider, D.M., Nelson, A., and Mooney, R. (2014). A synaptic and circuit basis for corollary discharge in the auditory cortex. Nature 513, 189–194. 10.1038/nature13724.

75. Liu, Y., Zhang, J., Jiang, Z., Qin, M., Xu, M., Zhang, S., and Ma, G. (2024). Organization of corticocortical and thalamocortical top-down inputs in the primary visual cortex. Nat Commun 15, 4495. 10.1038/s41467-024-48924-8.

76. Liu, Y., Bech, P., Tamura, K., Délez, L.T., Crochet, S., and Petersen, C.C. (2024). Cell class-specific long-range axonal projections of neurons in mouse whisker-related somatosensory cortices. Elife 13. 10.7554/eLife.97602.

77. Sievers, M., Motta, A., Schmidt, M., Yener, Y., Loomba, S., Song, K., Bruett, J., and Helmstaedter, M. (2024). Connectomic reconstruction of a cortical column. Preprint, 10.1101/2024.03.22.586254.

78. Akrami, A., Kopec, C.D., Diamond, M.E., and Brody, C.D. (2018). Posterior parietal cortex represents sensory history and mediates its effects on behaviour. Nature 554, 368–372. 10.1038/nature25510.

79. Mohan, H., Gallero-Salas, Y., Carta, S., Sacramento, J., Laurenczy, B., Sumanovski, L.T., de Kock, C.P.J., Helmchen, F., and Sachidhanandam, S. (2018). Sensory representation of an auditory cued tactile stimulus in the posterior parietal cortex of the mouse. Sci Rep 8, 7739. 10.1038/s41598-018-25891-x.

80. Rokach, R.O., Marmor, O., Levy, Y., and Gilad, A. (2025). ’What’ and ‘where’ brain-wide pathways are dominated by internal strategies. Preprint, 10.1101/2025.03.23.644791.

81. Benisty, H., Barson, D., Moberly, A.H., Lohani, S., Tang, L., Coifman, R.R., Crair, M.C., Mishne, G., Cardin, J.A., and Higley, M.J. (2024). Rapid fluctuations in functional connectivity of cortical networks encode spontaneous behavior. Nat Neurosci 27, 148–158. 10.1038/s41593-023-01498-y.

82. Lee, S., Kruglikov, I., Huang, Z.J., Fishell, G., and Rudy, B. (2013). A disinhibitory circuit mediates motor integration in the somatosensory cortex. Nat Neurosci 16, 1662–1670. 10.1038/nn.3544.

83. Raltschev, C., Kasavica, S., Leonardon, B., Nevian, T., and Sachidhanandam, S. (2025). Top-down modulation of sensory processing and mismatch in the mouse posterior parietal cortex. Nat Commun 16, 4240. 10.1038/s41467-025-58002-2.

84. Furutachi, S., Franklin, A.D., Aldea, A.M., Mrsic-Flogel, T.D., and Hofer, S.B. (2024). Cooperative thalamocortical circuit mechanism for sensory prediction errors. Nature 633, 398–406. 10.1038/s41586-024-07851-w.

85. Campagnola, L., Seeman, S.C., Chartrand, T., Kim, L., Hoggarth, A., Gamlin, C., Ito, S., Trinh, J., Davoudian, P., Radaelli, C., et al. (2022). Local connectivity and synaptic dynamics in mouse and human neocortex. Science (1979) 375. 10.1126/science.abj5861.

86. Lohani, S., Moberly, A.H., Benisty, H., Landa, B., Jing, M., Li, Y., Higley, M.J., and Cardin, J.A. (2022). Spatiotemporally heterogeneous coordination of cholinergic and neocortical activity. Nat Neurosci 25, 1706–1713. 10.1038/s41593-022-01202-6.

87. Sillito, A.M., and Kemp, J.A. (1983). Cholinergic modulation of the functional organization of the cat visual cortex. Brain Res 289, 143–155. 10.1016/0006-8993(83)90015-X.

88. Soma, S., Shimegi, S., Suematsu, N., Tamura, H., and Sato, H. (2013). Modulation-Specific and Laminar-Dependent Effects of Acetylcholine on Visual Responses in the Rat Primary Visual Cortex. PLoS One 8, e68430. 10.1371/journal.pone.0068430.

89. Yogesh, B., and Keller, G.B. (2024). Cholinergic input to mouse visual cortex signals a movement state and acutely enhances layer 5 responsiveness. Elife 12. 10.7554/eLife.89986.

90. Bethge, P., Carta, S., Lorenzo, D.A., Egolf, L., Goniotaki, D., Madisen, L., Voigt, F.F., Chen, J.L., Schneider, B., Ohkura, M., et al. (2017). An R-CaMP1.07 reporter mouse for cell-type-specific expression of a sensitive red fluorescent calcium indicator. PLoS One 12, e0179460. 10.1371/journal.pone.0179460.

91. Mayford, M., Bach, M.E., Huang, Y.-Y., Wang, L., Hawkins, R.D., and Kandel, E.R. (1996). Control of Memory Formation Through Regulated Expression of a CaMKII Transgene. Science (1979) 274, 1678–1683. 10.1126/science.274.5293.1678.

92. Harris, J.A., Hirokawa, K.E., Sorensen, S.A., Gu, H., Mills, M., Ng, L.L., Bohn, P., Mortrud, M., Ouellette, B., Kidney, J., et al. (2014). Anatomical characterization of Cre driver mice for neural circuit mapping and manipulation. Front Neural Circuits 8. 10.3389/fncir.2014.00076.

93. Guo, P.J., Kim, J., and Rubin, R. (2014). How video production affects student engagement. In Proceedings of the first ACM conference on Learning @ scale conference (ACM), pp. 41–50. 10.1145/2556325.2566239.

94. Abdelfattah, A., Allu, S.R., Campbell, R.E., Cheng, X., Cižmár, T., Costantini, I., Emiliani, V., Fomin-Thunemann, N., Gilad, A., Fernández Alfonso, T., et al. (2022). Neurophotonic Tools for Microscopic Measurements and Manipulation: Status Report. Neurophotonics 9. 10.1117/1.NPh.9.S1.013001.

95. Gilad, A. (2024). Wide-field imaging in behaving mice as a tool to study cognitive function. Neurophotonics 11. 10.1117/1.NPh.11.3.033404.

