## Supplementary figures for "Cortex-wide laminar dynamics diverge during learning"

**Supplementary material**

Supplementary Figure 1-6.

Supplementary Video 1-2.

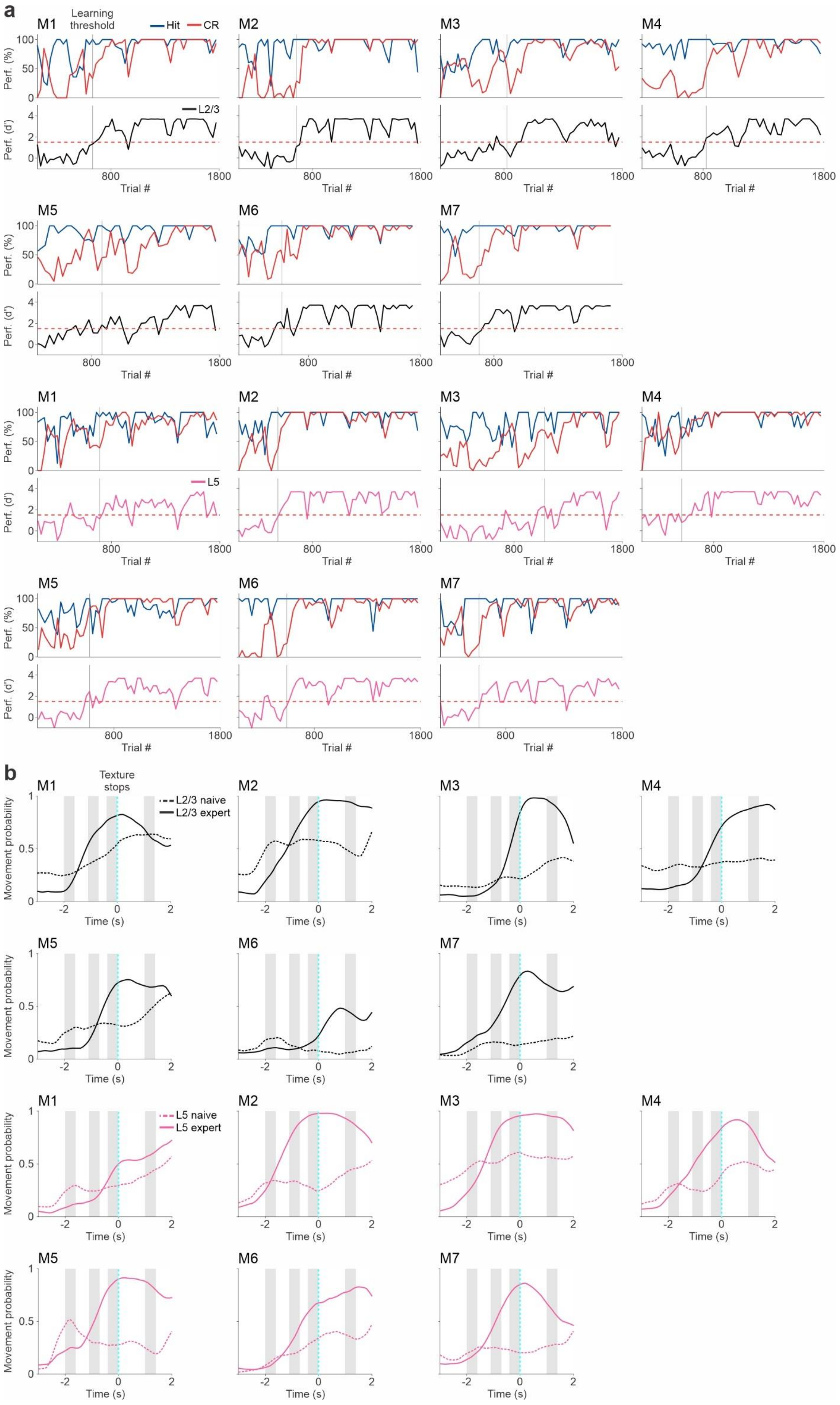
a
M1
Learning threshold
Hit
CR
M2
M3
M4
M5
M6
M7
Perf. (%)
Perf. (d')
L2/3
L5
800
1800
Trial #
b
Texture stops
L2/3 naive
L2/3 expert
L5 naive
L5 expert
Movement probability
Time (s)

**Supplementary Figure 1. Learning and movement changes during learning of individual mice. a** Top plot: Percentage (%) of Hit (blue) and CR trials (red) as a function of trials (30 trial bins) for each mouse separately. Bottom panel: Performance (d′) as a function of trial number feach mouse separately: L2/3 (black; n = 7) and L5 (pink; n = 7). Red dashed horizontal line indicates threshold for learning (d′ = 1.5) and gray vertical line indicates the learning threshold for each mouse. **b** Movement probability along the trial across all mice of L2/3 (black; n = 7) and L5 (pink; n = 7) mice during naïve (dashed lines) and expert (solid lines) cases. Gray bars indicate temporal periods.

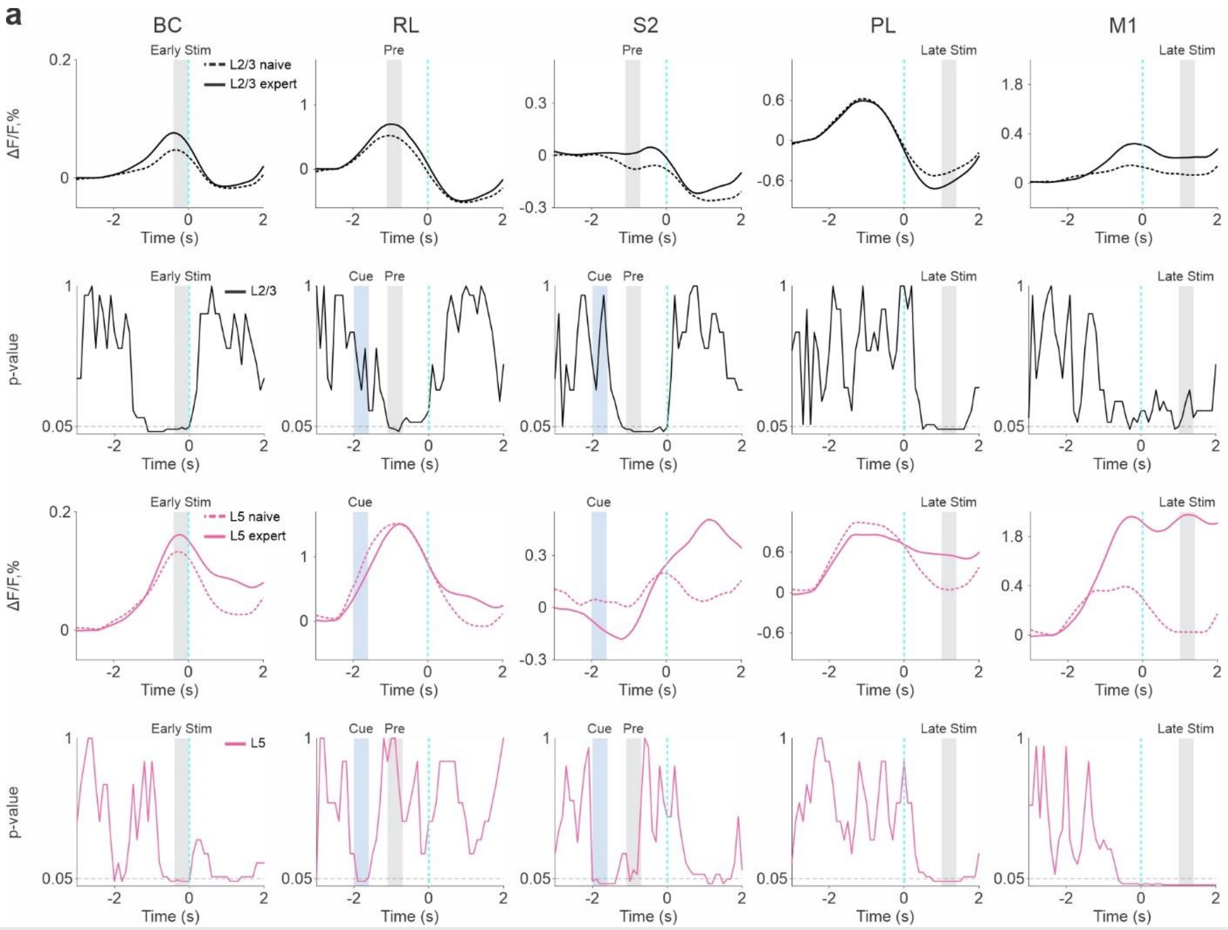


**Supplementary Figure 2. Frame-by-frame statistical comparison between naïve and expert for L2/3 and L5.** **a** Top plot: Temporal response (ΔF/F) along the trial for naïve (dashed) and expert (solid) cases plotted for each mouse separately in five areas of interest (from left to right): BC, RL, S2, PL, and M1 (as in Figure 2b). Expert (solid line) and naïve (dashed line). L2/3 in black and L5 in pink. Relevant time periods are marked by gray or blue colors. Bottom plot: Frame-by-frame p-value derived from a statistical comparison between naïve and expert trials (Rank-sum test) for each mouse separately (corresponding to the top plot). Significance threshold ($p = 0.05$) is marked in a gray dashed line.

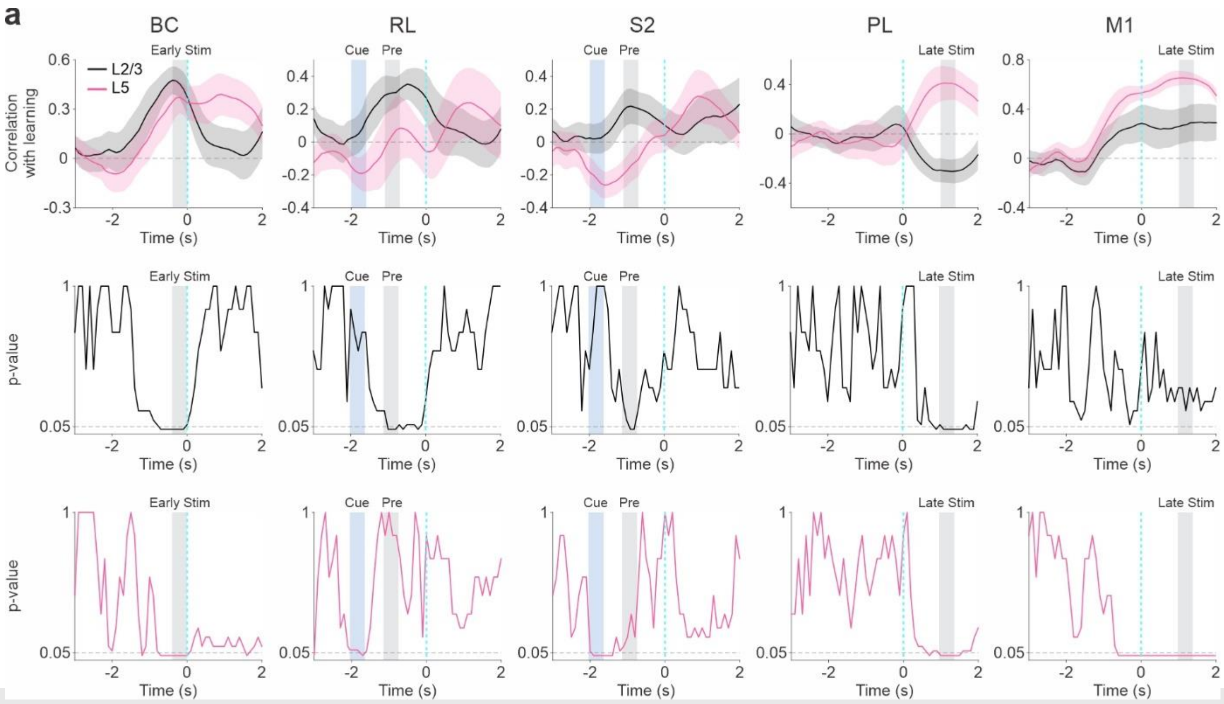


**Supplementary Figure 3. Frame-by-frame statistical analysis for correlation with learning. a** Top: Correlation with learning as a function of time for the 5 selected areas (from left to right) averaged across mice for L2/3 (black; n = 7) and L5 (pink; n = 7; As in Fig. 3c). Error shading indicates ± s.e.m. Time period for analysis window is marked by gray or blue. Bottom plots: Corresponding frame-by-frame p-value for a statistical test comparing correlation with learning values with 0 (Signed-rank test; n = 7 mice for each layer. L/23 in black and L5 in pink. Significance threshold (p = 0.05) is marked in a gray dashed line.

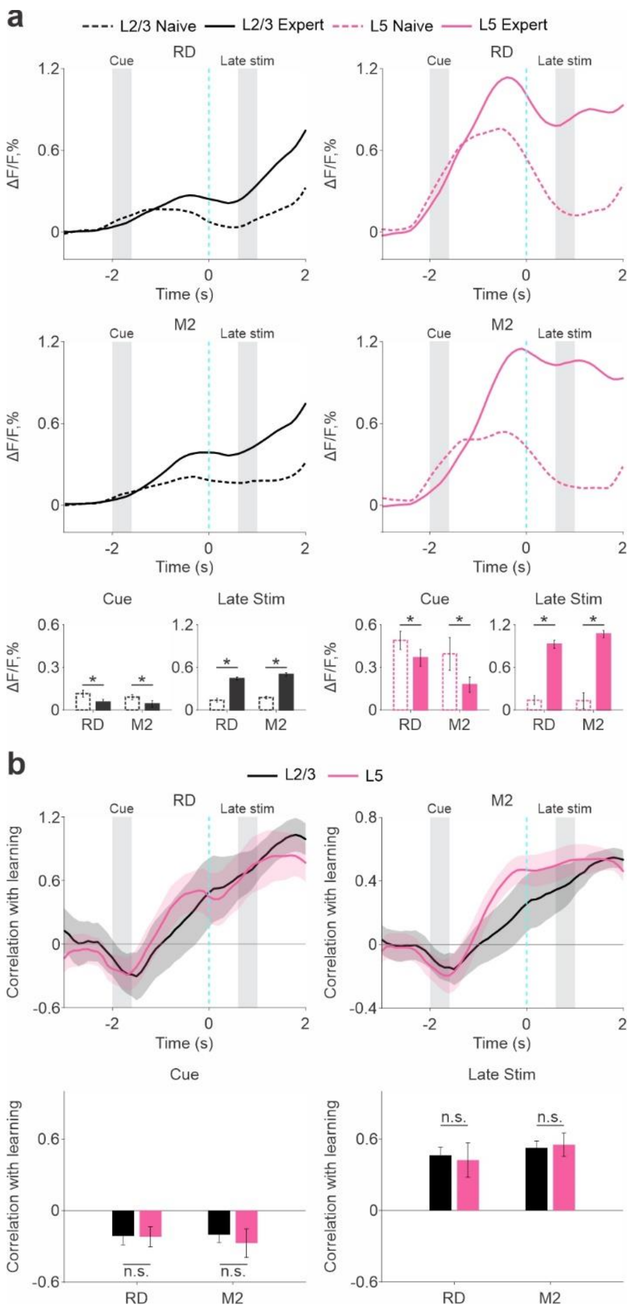


**Supplementary Figure 4. M2 and RD display similar learning-related parameters in L2/3 and L5. a** Top: Temporal response (ΔF/F) along the trial for RD (top) and M2 (middle) averaged across mice of L2/3 (left, black; n = 7) and L5 (right, pink; n = 7) for expert (solid line) and naïve (dashed line) cases. Mean response for RD and M2 (bottom) during cue (left) and late stim (right) in expert (straight line) and naïve (dashed line) cases. Error bars are s.e.m. across mice (n = 7 for each layer). Lines depict individual mice. *$p < 0.05$, n.s. not significant, Rank-sum test. **b** Top: correlation with learning as a function of time for areas RD (left) and M2 (right) averaged across mice for L2/3 (black; n = 7) and L5 (pink; n = 7). Error shading indicates ± s.e.m. Time period for analysis window is marked by gray or blue. Bottom: Correlation with learning for each area during cue (left) and late-stim (right) indicated at the top. Error bars are s.e.m. across mice. Gray dots depict individual mice. n.s. not significant, Rank-sum test. In general RD and M2 parameters are similar across layers.

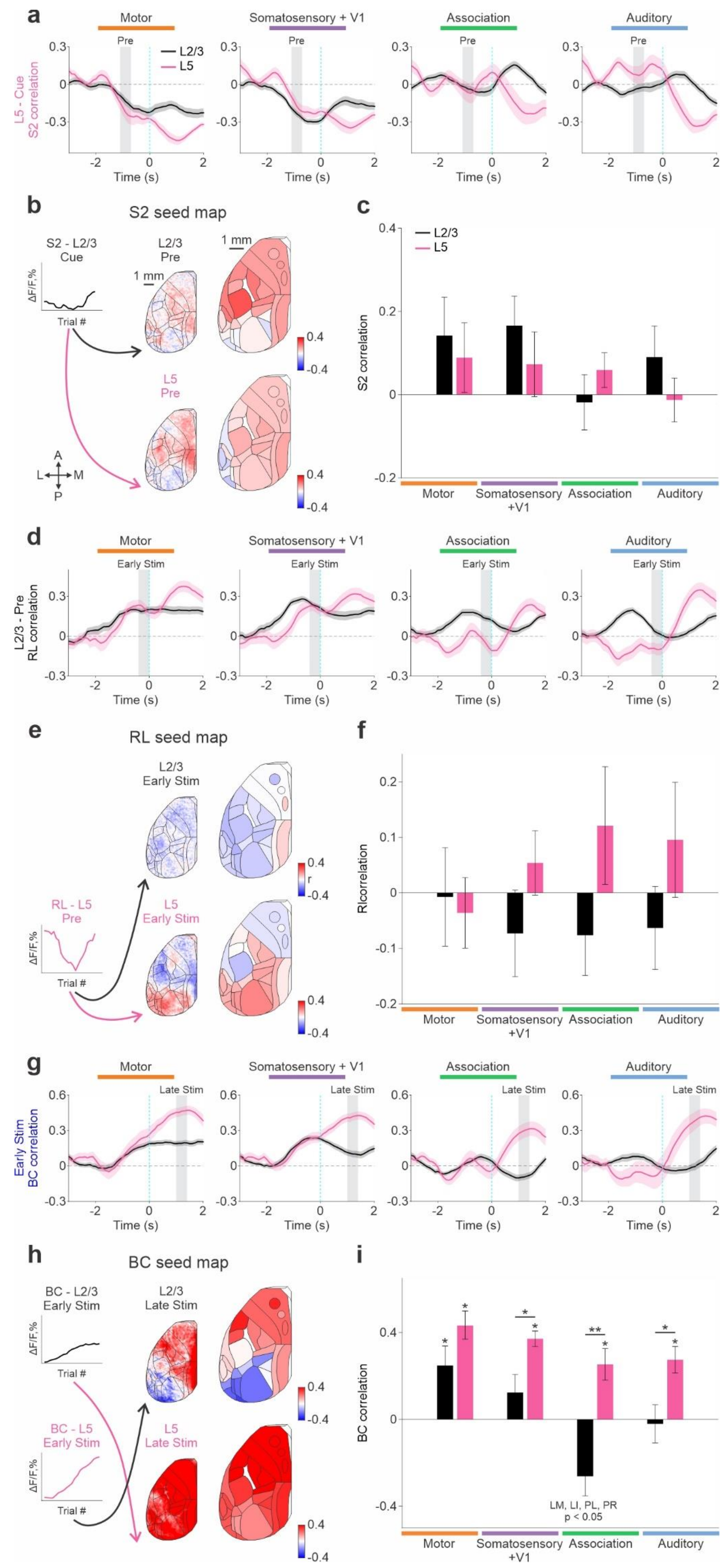
a
Motor
Somatosensory + V1
Association
Auditory
Pre
L2/3
L5
L5 - Cue
S2 correlation
Time (s)
b
S2 seed map
S2 - L2/3
Cue
L2/3
Pre
1 mm
ΔF/F,%
Trial #
L5
Pre
A
L
M
P
c
L2/3
L5
S2 correlation
Motor
Somatosensory
+V1
Association
Auditory
d
Early Stim
L2/3 - Pre
RL correlation
Time (s)
e
RL seed map
L2/3
Early Stim
RL - L5
Pre
L5
Early Stim
r
f
Rlcorrelation
g
Late Stim
Early Stim
BC correlation
h
BC seed map
BC - L2/3
Early Stim
L2/3
Late Stim
BC - L5
Early Stim
L5
Late Stim
i
BC correlation
LM, LI, PL, PR
p < 0.05

**Supplementary Figure 5. ROI seed correlations in time and additional maps. a** Seed correlation between L5 S2 learning curve during the cue-period and the learning curves in each time frame, grouped for Motor, Somatosensory + V1, Association and Auditory (left to right). L2/3 in black and L5 in pink. Error shading indicates ± s.e.m across mice (n = 7). Time period for analysis window is marked by gray color. **b** ROI seed map complementing Figure 6c-d, by taking the learning curve in S2 L2/3 during the cue-period. The small map is an example mouse, and the big map is the average across mice (n = 7). **c** Seed correlation values grouped into Motor, Somatosensory + V1, Association and Auditory for L2/3 and L5. **d,e,f** same as a,c,d but the seed is RL in L5 during the pre-period and the map is L2/3 or L5 during the early-stim period (complementing Figure 6e-f). **g,h,i** same as a,c,d but the seed area is BC either in L2/3 or L5 during the early-stim period and the map is either L2/3 or L5 during the late-stim period (complementing Figure 6g-h). $^{*}p < 0.05$, $^{**}p < 0.005$, Signed-rank test within layer, Rank-sum test across layers.

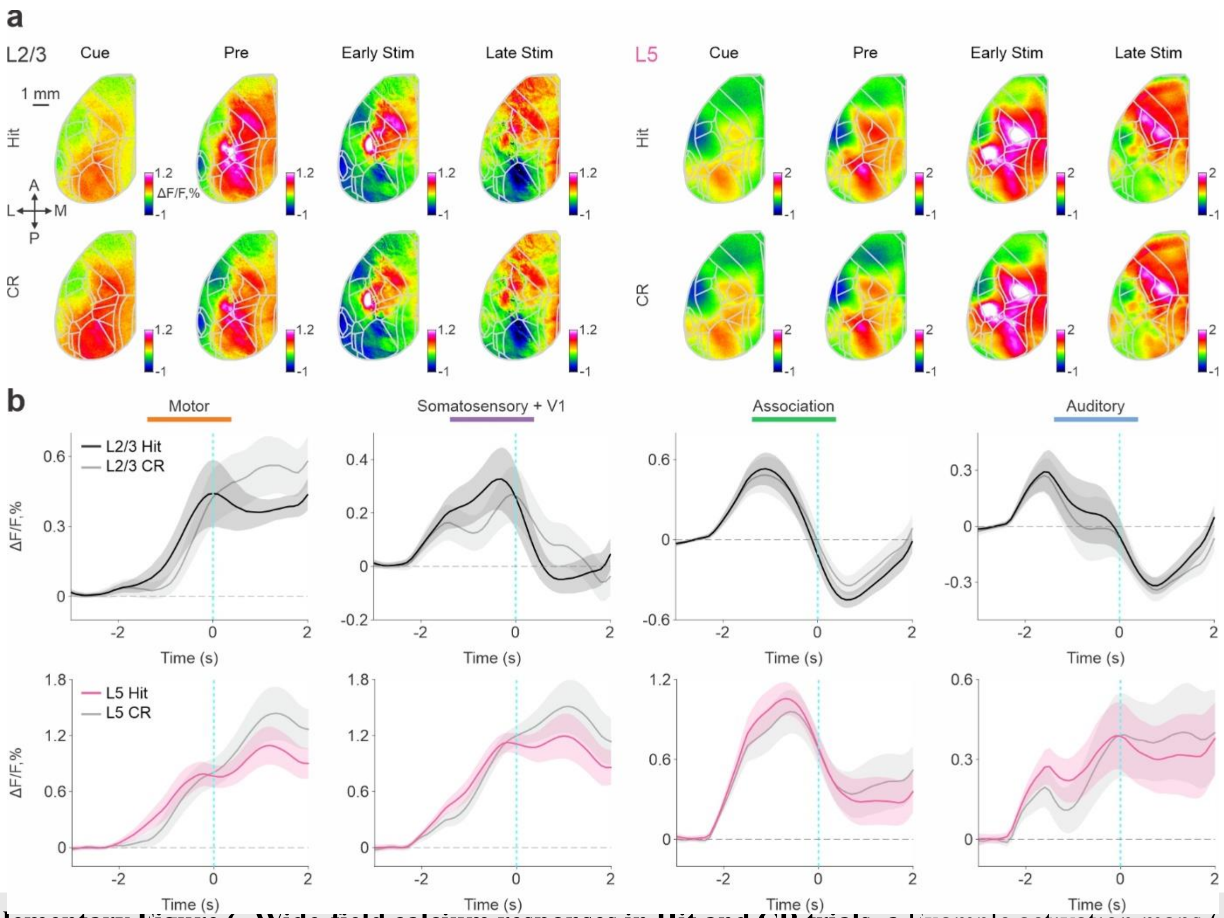


**Supplementary Figure 6. Wide-field calcium responses in Hit and CR trials. a** Example activation maps (ΔF/F) in one mouse for L2/3 (left) and L5 (right) averaged during cue, pre, early-stim and late-stim periods in Hit (top) and CR (bottom) expert case. Color scale bar indicates min/max of percent ΔF/F. Overlay of areas in gray for all maps. Scale bar is 1 mm. **b** Temporal response along the trial for each grouped areas Motor, Somatosensory + V1, Association and Auditory (left to right) averaged across L2/3 (black, top) and L5 (pink, bottom) mice (n = 7 for each mouse). Error shading indicates s.e.m.

**Supplementary Video 1. Cortex-wide laminar differences during learning as a function of time.** Example activation maps (ΔF/F) as a function of time of one L2/3 (left) and L5 (right) mouse during naïve (top) and expert (bottom) phases. Color denotes ΔF/F of percent. Color scale bar for L2/3 min -1 max 1.2 and L5 min -1 max 2.

**Supplementary Video 2. Learning maps as a function of time.** Example learning maps (r) as a function of time of one L2/3 (left) and L5 (right) mouse. Color denotes r. Color scale bar for L2/3 and L5 min -0.5 max 0.7.
